# Herbarium specimens reveal a constrained seasonal climate niche despite diverged annual climates across a wildflower clade

**DOI:** 10.1101/2025.02.28.640808

**Authors:** Megan Bontrager, Samantha J. Worthy, N. Ivalú Cacho, Laura Leventhal, Julin N. Maloof, Jennifer R. Gremer, Johanna Schmitt, Sharon Y. Strauss

**Affiliations:** Department of Evolution and Ecology, UC Davis, Davis, CA, 95616; Center for Population Biology, UC Davis, Davis, CA, 95616; Department of Ecology and Evolutionary Biology, University of Toronto, Toronto, Ontario, Canada, M5S 3B2; School of Biological Sciences, University of Nebraska, Lincoln, NE, 68520; Instituto de Biología, Universidad Nacional Autónoma de México, Ciudad de México, México; Department of Plant Biology, Carnegie Institution for Science, Stanford, CA, 94305; Department of Biology, Stanford University, Stanford, CA, 94305; Department of Integrative Biology, University of California Berkeley, Berkeley, CA, 94720; Department of Plant Biology, UC Davis, Davis, CA, USA, 95616

**Keywords:** climate niche evolution, Brassicaceae, seasonal niche, climate change, phenology, germination, microrefugia, niche construction

## Abstract

Quantifying species’ niches across a clade reveals how environmental tolerances evolve, and offers insights into present and future distributions. We use herbarium specimens to explore climate niche evolution across 14 annual species of the Streptanthus (s.l.) clade (Brassicaceae), which originated in deserts and diversified into cooler, moister areas. To understand how climate niches evolved, we used historical climate records to estimate each species’ 1) classic annual climate niche, averaged over specimen collection sites; 2) growing season niche, from estimated specimen germination date to collection date, averaged across specimens (specimen-specific niche); and 3) standardized seasonal niche based on average growing seasons of all species (clade-seasonal niche). In addition to estimating how phenological variation maps onto climate niche evolution, we explored how spatial refugia shape the climate experienced by species by 1) analyzing how field soil texture changes relative to the climate space that species occupy and 2) comparing soil water holding capacity from each specimen locality to that of surrounding areas. Specimen-specific niches exhibited less clade-wide variation in climatic water deficit than did annual or clade-seasonal niches, and specimen-specific temperature niches showed no phylogenetic signal, in contrast to annual and clade-seasonal temperature niches. Species occupying cooler regions tracked hotter and drier climates by growing later into the summer, and by inhabiting refugia on drought-prone soils.

These results underscore how phenological shifts, spatial refugia, and germination timing shape “lived” climate. Despite occupying a large range of annual climates, we found these species are constrained in the conditions under which they thrive.

**Significance statement:** Here, we show that closely related species that appear to have diverged in their climate niches based on highly differentiated annual conditions are actually tracking very similar seasonal climate conditions. This climate similarity, estimated from collection locations and associated weather data of nearly 2000 herbarium specimens, is achieved through the evolution of germination timing and plant flowering phenology, as well as establishment in spatial refugia with suitable microclimates. Our results highlight how occupation of a subset of seasonal conditions can be conserved across diverse species, resulting in less climate niche evolution than expected. Restricted climate niches imply that species may be less adaptable than we expect based on annual climates, and have implications for conservation, management and persistence under hotter climates.

## Introduction

The geographic distributions of species within a clade can reflect how environmental tolerances and niches have evolved, and may reveal evolutionary constraints on adaptation to changing climates. Insights may be improved by focusing on niche parameters during the temporal window when species are active. Organisms can influence their own growing conditions by adjusting their phenology to restrict activity and life cycle events to a narrower range of environmental conditions than those available to them (1–4). For example, mammals that hibernate during unfavorable winter conditions can occupy environments that would otherwise be too cold in winter (5). Similarly, germination cues used by plants allow them to select seasonal windows of environmental conditions that are favorable for future growth and reproduction (3, 6, 7).

In addition to selecting periods of time that facilitate growth and reproduction, organisms also sample subsets of space that represent suitable conditions (1, 8, 9). This ability might be obvious in some taxa, for example, many animals can move to occupy favorable temperatures by migrating seasonally (1, 10). However, sessile organisms can also occupy favorable microhabitats in areas of otherwise unsuitable habitat or climate through differential establishment in suitable microrefugia (8, 11–14) or by attracting dispersal agents that selectively deposit seeds in suitable habitats (8). Fine-scale spatial variation in the abiotic niche for plants comes from factors including geographic slope and aspect, as well as biotic and abiotic soil attributes (13–16). For instance, tree species adapted to hotter and drier habitats are able to survive on south-facing slopes of less than a kilometer within a matrix of less suitable cooler and wetter habitats (12). Similarly, temperate species that have dispersed southward during glaciations survive in gullies characterized by cooler temperatures in otherwise tropical areas (17). Ultimately, sessile species live in subsets of available conditions by constructing niches that are temporally narrower than the range of annual conditions, and by occupying subsets of suitable spatial habitat.

Museum collections offer valuable tools for understanding how species sample available environmental conditions, as these collections are temporally and spatially explicit and often cover broad temporal ranges and geographic space (18). Herbarium specimens are particularly useful as most plants spend their entire lives fixed at the same location. Their label data, coupled with data from climatic and geological databases, can provide detailed estimates of the seasonal growing conditions experienced by the collected individual (represented by the specimen), thereby opening new doors to understanding current distributions as well as trait and niche evolution (4, 19–23). For annual organisms, reconstructing the seasonal climate niche, which only considers the time window when an individual was alive or non-dormant, may allow for a better estimate of the climate factors determining species distributions in space and time than estimates based on annual climate data (4, 22, 24). This level of detail can enhance our ability to characterize niche evolution across clades.

Here, we study the evolution of the seasonal niche in a clade of primarily annual plant species native to the California Floristic Province (CFP), a biodiversity hotspot encompassing most of California, eastern Nevada, southern Oregon, and northern Baja California, Mexico (https://ucjeps.berkeley.edu/eflora/geography.html). The Mediterranean climate that dominates in the CFP consists of a wet season that begins in fall and ends in spring, with no appreciable precipitation during the 3-4 months of hot, dry summer. This Mediterranean climate pattern is relatively recent, originating 2-5 million years ago, and its onset facilitated northward range expansion of southwest desert-adapted species (25). In many groups, northward migration was accompanied by speciation, and these groups constitute what is termed the Madro-Tertiary flora, which comprises 14% of the present day flora of the CFP (25). If climate niches evolved as species diversified from the southwestern and western deserts to northern, more mesic habitats, then we would expect the niches of species that live further north to reflect the cooler, wetter growing seasons in these areas. In contrast, if the niche has remained relatively constant as species diversified, perhaps indicating some evolutionary constraint, we would expect the species to evolve different life cycle timing and habitat selection cues such that they maintain their preferred climate conditions, regardless of their geographic location.

One example of the Madro-Tertiary flora is the jewelflower (*Streptanthus* s.l.) clade in the family Brassicaceae, composed primarily of annual species (26–28). Using an existing phylogeny (27), we assess how the climate niche evolved across the clade as it speciated from the southwest desert to northern or high elevation habitats. We ask: 1) do species’ climate niches diverge with speciation into novel annual climates, and 2) to what degree do species alter their growing niche conditions via phenology of germination and senescence, or by occupying a subset of available microhabitats? We use spatially and temporally specific information from herbarium specimens, field data, and data from climate and geological databases to characterize the conditions under which these species grow and reproduce. Using precipitation records and prior experimental data on germination cues, we estimate the germination month for each specimen. Coupled with the collection date, we then estimate the “lived” climate experienced by each specimen (referred to as the “specimen-specific” climate niche). These values could then be compared to those calculated across annual and standardized seasonal (“clade-seasonal”) time windows calculated to represent the average growing season across the clade. This approach allows us to consider mechanisms of specifically timed phenology that determine growing conditions, also known as niche construction (1, 2, 29), as we explore the extent of niche evolution with speciation into new geographic regions.

## Results

### Phenological variation across the clade

The 14 focal annual species varied in which months they grew, with many of the lower latitude southern species starting their lives (and growing seasons) later and completing them before northern species (Figure 1C, S2, S3). Lower latitude species likely start the growing season slightly later due to substantial rainfall arriving later in these regions (Figure S2, S3A)– rain plays a key role as cue for germination in this clade (30). In general, lower-latitude species have shorter estimated lifespans than higher-latitude ones (Figure S2, S3C). This result is consistent with experimental work in this system, in which low-latitude *Caulanthus* species did not extend their flowering time, even under abundant water availability (31).

**Figure 1.**
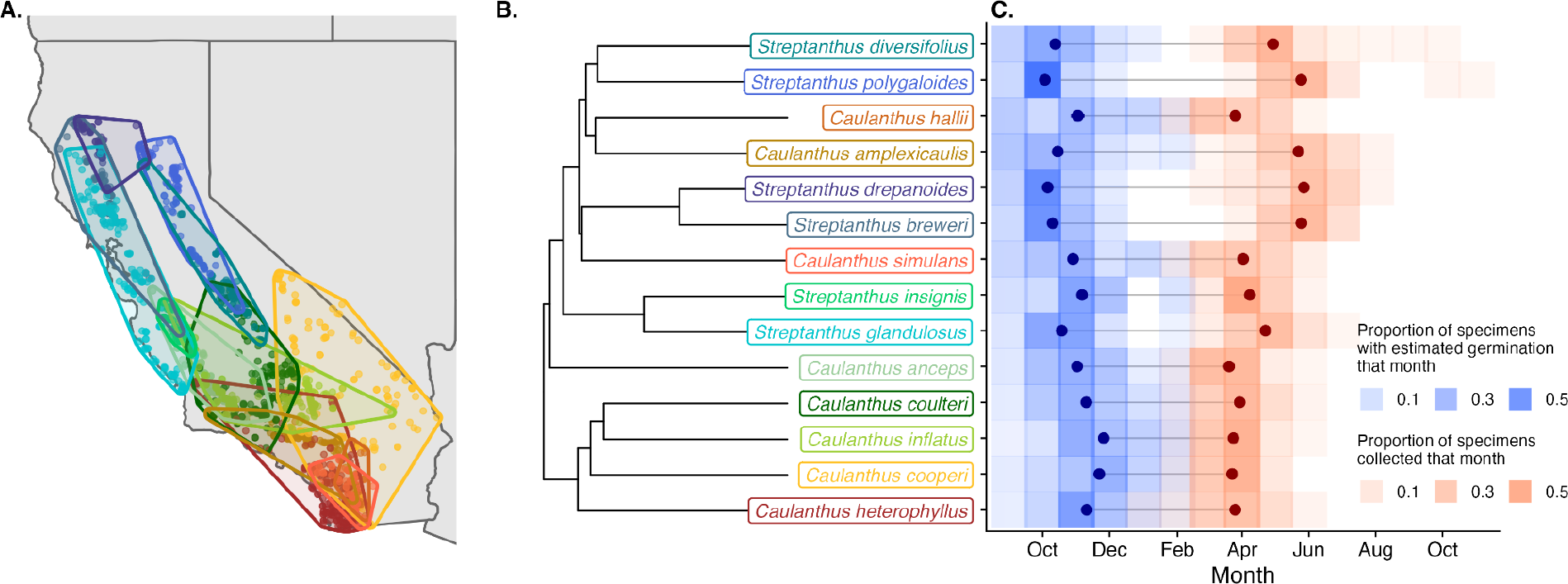
Geographic distributions (A.), phylogenetic relationships (B.), and specimen-specific seasonal windows (C.) of the 14 annual species from the major subclades in the larger Streptanthus I clade included in this study. Map polygons in (A.) represent locations of herbarium specimens for each species that were used in this study (most specimens were located in California, USA). The phylogeny in (B.) is from (27), trimmed to our focal species. The color of the species’ points and polygons on the map matches the color of the species’ names on the phylogeny. Shaded tiles in (C.) represent the relative proportion of specimens that were estimated to have germinated (blue) or that were collected (red) in a given month for each species. Point ranges in (C.) represent the average germination month estimate (dark blue) and collection date (dark red) for each species.

### Evolution of the annual and seasonal climate niche

Phylogenetic niche conservatism (PNC) is detected when closely related species retain aspects of their niches over time and are more similar than would be expected by Brownian motion (neutral) evolution (32–34). One can also ask if trait evolution is consistent with Brownian motion evolution that is bounded by character states set by selection, physical processes or both (e.g. (34, 35)), in our case we used clade-wide climate variable minimum and maximum values. We also considered the possibility of stabilizing selection on climate niche trait evolution, estimated with an Ornstein–Uhlenbeck (OU) process that includes constraints on the variance that a trait can attain in a clade (e.g. (36, 37)) and tested which of these models of evolution best fit our climate niche evolution data. Annual climate niches were very diverged among species (Figures S4-S6), as might be expected for a group of species that span 3000 m in elevation, nearly 8 degrees of latitude, and occur in diverse habitats (Figures 1, S1). To look at the evolution of the climate niche in more detail, we compared climatic water deficit (CWD), temperature, and precipitation niches across annual, clade-seasonal (Oct-April), and specimen-specific seasonal climate intervals, and examined how variation in these niche axes is distributed across the phylogeny.

Evolution of the annual CWD niche was best fit by a bounded Brownian motion (BBM) model of evolution. We then asked whether evolution was more phylogenetically conserved than expected under a traditional unbounded Brownian model (K significantly greater than 1) (38) (Table S1, S2, Figure S4), potentially indicating phylogenetic niche conservatism (33). Annual temperature and precipitation evolution also showed significant phylogenetic signal (Table S1, Figures S5 and S6), with temperature evolution best fit by a Brownian motion model and precipitation by a bounded Brownian motion model (Table S2). Phylogenetic structure in annual climate stems in part from climate variation reflecting clade origins, with the “true” *Caulanthus* subclade (*C. coulteri, C. heterophyllus, C. inflatus,* and *C. cooperi*) occupying hotter, drier climates than most other species (Table S1, Figures 2, S4-S6).

**Figure 2.**
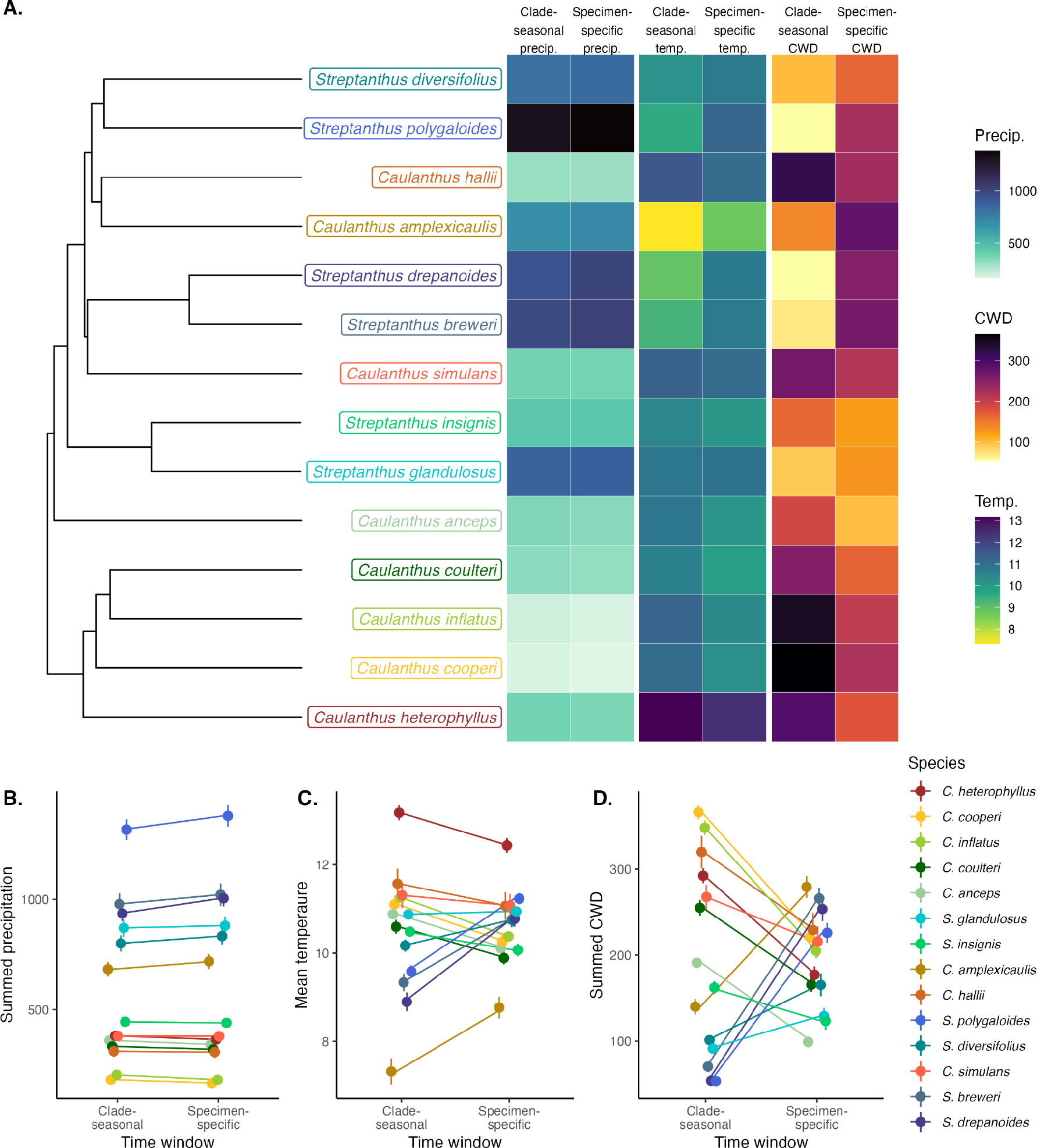
(A.) Visual comparisons of the specimen-specific seasonal climate niche (climate spanning germination month to collection month, averaged over specimens) and average clade-wide seasonal climate niche (Oct-Apr of the collection year) plotted alongside the phylogeny. The three climate variables are total precipitation (mm), average temperature (°C), and total climate water deficit (CWD). Panels (B.), (C.), and (D.) compare average clade-wide and specimen-specific niches for each of the three climate variables; points represent species’ means and error bars represent one standard error. The variances (of species’ means) of the average clade-wide and specimen-specific time windows were not different for precipitation (B.) or temperature (C.), but did significantly differ for CWD (D.), with lower variance when CWD was summarized over specimen-specific time windows.

Our estimates of the climate niche based on the clade-seasonal time window (April-October) were highly correlated with the annual climate niche (r >= 0.93, Table 1, Figures S4-S7); thus, seasonal subsampling without phenological variation among species and specimens does not strongly alter the conditions that species occupy relative to each other.

**Table 1.** Correlations of phylogenetically independent contrasts between climate variables summarized across each of the three time windows. Bold text indicates that the correlation is significant (p < 0.05). Also presented are 95% confidence intervals.

| Climate variable | Annual vs. clade-seasonal | Annual vs. specimen-specific | Clade-seasonal vs. specimen-specific |
| --- | --- | --- | --- |
| Temperature | **$r = 0.931$ [0.781-0.980]** | **$r = 0.609$ [0.087-0.869]** | **$r = 0.761$ [0.362-0.924]** |
| Precipitation | **$r = 0.999$ [0.997-1.000]** | **$r = 1.000$ [0.999-1.000]** | **$r = 0.999$ [0.998-1.000]** |
| Climatic water deficit | **$r = 0.942$ [0.814-0.983]** | $r = -0.220$ [-0.688-0.376] | $r = 0.05$ [-0.512-0.588] |

Overall, the clade-seasonal time window had lower temperature and CWD than the annual time window because it does not include the warmest and driest months of the year (Figures S4, S5). All time windows encompassed the vast majority of annual precipitation, so precipitation values were highly correlated and similar across all three time windows (r >= 0.99, Table 1, Figure S6, S7D-F). Patterns of phylogenetic signal for the clade-seasonal niche mirrored those of the annual niche—evolution of the clade-seasonal CWD niche was more constrained than expected under Brownian evolution (K significantly > 1; Table S1), consistent with phylogenetically constrained evolution, with bounded and unbounded Brownian evolution as the best models (Table S2). Temperature and precipitation showed significant signal consistent with bounded Brownian evolution (Table S1, S2). While all these models help us to understand trait divergence in relation to phylogenetic divergence, it is much harder to infer phylogenetic niche conservatism from these analyses (32, 39). Below we explore patterns of trait variation that may shed further light on niche evolution and constraint.

If species are retaining aspects of their ancestral climate niche, then we would expect them to be tracking similar growing conditions even as they occupy different geographic regions and as they diverge. Consistent with this expectation, specimen-specific time windows—which span the month of first rain in each specimen’s collection site until the collection month of the specimen—comprise different temperature and CWD conditions than annual and clade-seasonal niches (Figure 2). Specimen-specific estimates of the CWD niche were not correlated with estimates of the CWD niche at the clade-seasonal time scale (r = 0.05, Table 1, Figure S7H) or the annual time scale (r = -0.22, Table 1, Figure S7I), indicating that local phenology is substantially changing the relative CWD conditions these species experience. In addition, we found significantly lower variance in CWD in specimen-specific relative to clade-seasonal windows (variance 3043 vs. 12734; Levene’s test F = 10.12, p = 0.0038, Figure 2D) and a narrower range of CWD values calculated over specimen-specific (range: 99-279) vs. clade-seasonal average time windows (range: 54-366). These results demonstrate that small changes in the timing of the growing season can shape the climate that these species experience (Figure 2), and suggest constraints in climate adaptation indicated by the restricted range of CWD across species in the clade-seasonal niche. We interpret this markedly reduced variation in the specimen-specific CWD niche across the clade as an indicator of evolutionary constraint and phylogenetic niche conservatism.

The specimen-specific temperature niche also exhibited reduced variation among species. Although the correlation between temperature estimated in specimen-specific and clade-seasonal time windows was fairly high (r = 0.76; Table 1), phenological variation among species and individuals somewhat alters the estimated thermal niches that species occupy relative to each other (Figure 2A,C, S6). Specimen-specific temperature niches exhibited a narrower range of values than clade-seasonal niches (clade-seasonal range: 7.3-13.2 (5.9°C), specimen-specific range: 8.8-12.4 (3.6°C). In addition, the temperature niche in specimen-specific time windows showed lower variance among species (variance = 0.69) than did estimates in the clade-seasonal time window (variance = 1.93, Figure 2A,C), though these variances were not significantly different (Levene’s test F = 1.77, p = 0.20). Unlike other aspects of the climate niche, the specimen-specific temperature niche showed no phylogenetic signal (Table S1). In other words, close relatives do not tend to have more similar thermal niches once specimen-specific timing is taken into account.

Specimen-specific climate niches also bucked latitudinal climate trends. Surprisingly, specimen-specific CWD in some more northern species (*S. drepanoides, S. breweri, S. polygaloides)* was greater than that exhibited by species with southern distributions (e.g., *C. heterophyllus, C. simulans, C. inflatus*) whose annual and clade-seasonal climates are hotter and drier (much greater annual CWD) than these northern species (Figure 2A,D). These northern species are occupying a seasonal niche with relatively higher CWD via longer lifespans extending into the hotter, drier months (Figures 1, S2, S3), while more southern species in annually hotter climates are restricting their growing season to cooler months (compare annual, clade-seasonal and specimen-specific CWD in Figures 2A,C, S4). This pattern is supportedby a significant relationship between mean latitude and the difference between specimen-specific and clade-seasonal CWD (PGLS slope = 42.7, p = 0.003; Figure 4). Species from lower latitudes track cooler temperatures relative to clade-seasonal niches while those at higher latitudes track relatively warmer climates, again suggesting evolutionary constraint in thermal tolerances (Figure 4). Species did not have significantly lower variances in specimen-specific vs. clade-seasonal temperature niches (p = 0.19; Figure 2C), nor in clade-seasonal and specimen-specific precipitation (Levene’s test F = 0.055, p = 0.82; Figure 2B), as all time windows sample all months with substantial precipitation.

**Figure 3.**
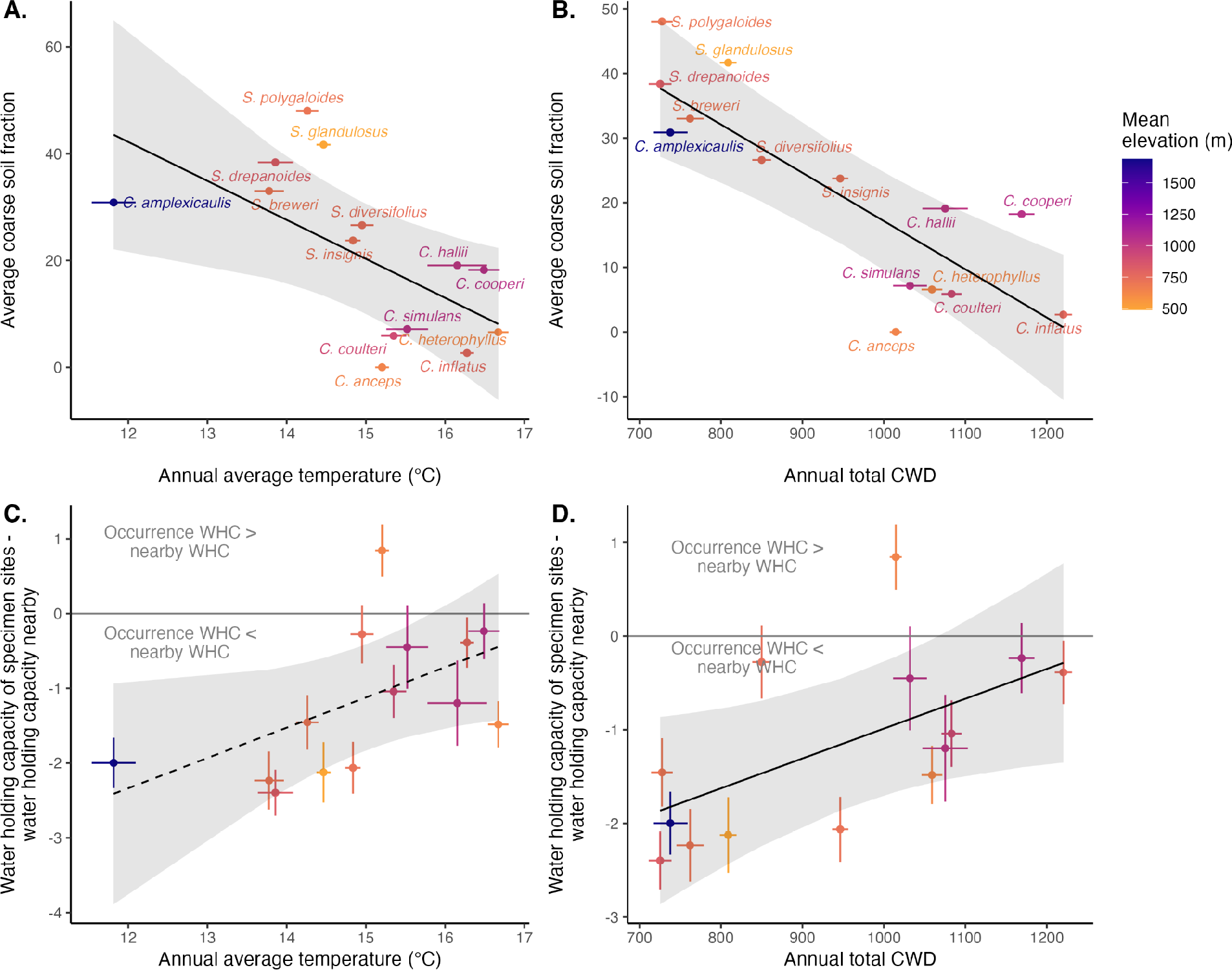
(A.) Species occurring in cooler areas (lower annual average temperature) occur in soils with greater fractions of coarse material. Line is the regression slope from a PGLS model (slope = -7.3, p = 0.019). Coarse soil fraction was measured from field soil sampled from the rhizosphere of *Streptanthus* or *Caulanthus* individuals in an average of 4.3 sites per species. (B.) Species occurring in more mesic areas (lower annual climatic water deficit; CWD) occur in soils with greater fractions of coarse material. Line is the regression slope from a PGLS model (slope = -0.075, p < 0.001). Point colors in (A.) and (B.) represent elevation; species in our analyses had mean elevations between 490 and 1050 m, except for *C. amplexicaulis* which had a mean elevation of 1690 m. (C.) Species occurring in cooler areas tend to occupy grid cells that have lower water holding capacity (WHC) than the surrounding area, while those in warmer areas occupy sites more similar to the surrounding areas, but this trend is only marginally significant (slope from a PGLS model = 0.40, p = 0.051). (D.) Species occurring in more mesic areas (low CWD) occupy grid cells that have lower water holding capacity than the surrounding area, while those in more arid areas (high CWD) occupy sites more similar to the surrounding areas. Line is the regression slope from a PGLS model (slope = 0.0032, p = 0.042). In all panels, shaded bands represent 95% confidence intervals generated using the function ggpredict() from the package ggeffects (78) and error bars on points represent standard errors.

**Figure 4.**
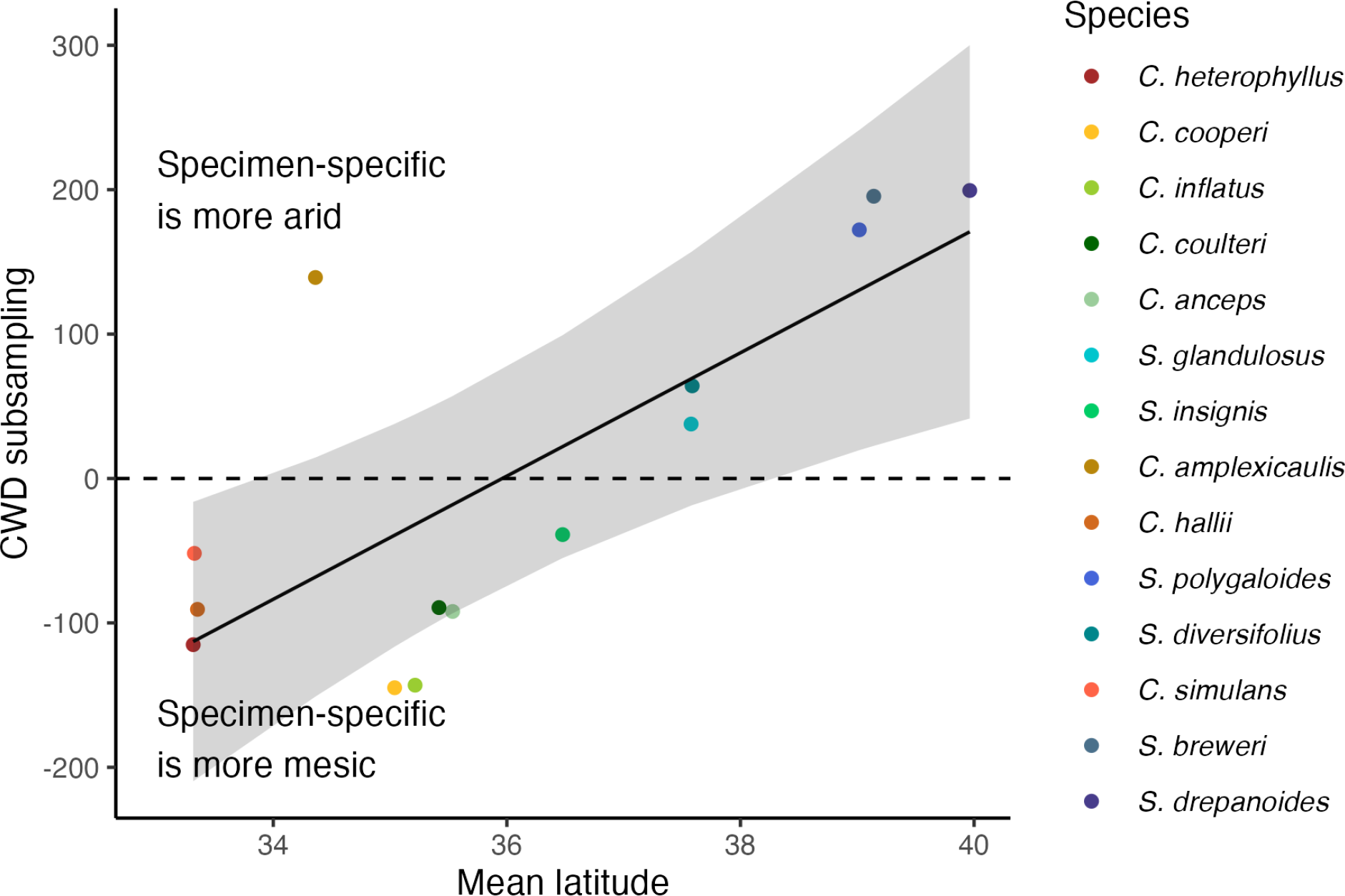
Species living at latitudes that vary in CWD adjust their phenology to track similar conditions. Species track more arid CWD when they live in northern areas and track more moist CWD when they live in southern areas. We detect this by taking the difference between CWD summed over specimen-specific time windows for each species and CWD summed over the standard clade-seasonal time window (Oct-April for all species). Positive values indicate that CWD calculated based on individual specimen phenology is higher (e.g., lived conditions are more arid) than CWD in a standard time window. Negative values indicate that specimen-specific phenology results in more mesic lived conditions. The regression line is from a PGLS model (slope = 42.7, p = 0.003). The shaded band represents the 95% confidence interval generated using the function ggpredict() from the package ggeffects (78). See also Figure 2D.

### Refugia driven by soil characteristics and water availability across the clade

Aside from constructing niches via the timing of when and how long to grow, species may also occupy spatial refugia in more permeable soils and hotter geographic aspects that result in lower water availability typical of ancestral desert habitats. Using data from soil collected at field sites directly under plants, we find that species that occupy cooler areas grow in substrates with higher fractions of coarse material relative to members of the clade living in warmer areas (Figure 3A; PGLS slope = -7.3, p = 0.019). Similarly, we find that species that live in more mesic, low CWD climates occupy soils with greater coarse fractions (Figure 3B; slope = -0.075, p < 0.001). Most species (13 of 14) occupy soils with relatively lower water holding capacity than the surrounding area (Sign test p = 0.0018, Figure S8). In particular, species that occupy cooler and more mesic areas occur in soils with lower water holding capacity than the surrounding area (e.g., Figure S9), while species that occur in warmer and drier areas occupy soils that do not differ as strongly in water holding capacity from the surrounding area (Figure 3C,D, PGLS temperature slope = 0.40, p = 0.051; CWD slope = 0.0032, p = 0.042). Thus differential establishment in suitable spatial microrefugia also contributes to the maintenance of more similar niches than expected based on latitudinal climate gradients.

## Discussion

This study shows that the timing of germination and reproduction, as well as differential establishment in soils with lower water availability, results in much less realized clade-wide climate niche evolution than expected given the widely divergent environments occupied by annual species in the Streptanthus *s.l.* clade. We gained this understanding of constraint in niche evolution by combining herbarium specimen data, climate and soil data, and our knowledge of germination behavior in this clade, which allowed us to estimate specimens’ growing seasons. Had we relied only on the annual niche metrics that are typically used in species distribution models and ecological niche models (reviewed in (24)) or even a standard seasonal niche for the clade (like the clade-seasonal niche), we might have concluded that species in this clade exhibit varied climate niches and underwent extensive climate niche evolution as they moved to cooler, moister habitats from ancestrally dry ones. However, we show that phenological shifts, coupled with differential establishment (or directed dispersal (8, 40)) across microhabitats, allows these species to track very similar realized climates, especially temperature and climate water deficit. For example, the hotter, drier seasonal niche of species in more mesic habitats is achieved by occupying dry microsites and by growing for a longer time into hot, dry seasons. This longer season may reflect common developmental requirements across the clade, e.g., a longer season may allow species in cooler areas to accumulate the growing degree-days necessary to complete their life cycles. The low variability in growing season CWD and similar trends in temperature experienced by these species across variable environments suggests that there may be strong evolutionary constraints on temperature and moisture tolerance in this clade.

Several other studies have suggested the importance of considering only the months in which species are seasonally active to estimate niche properties (4, 10, 23, 41). In a recent survey, Zurell and colleagues (24) found that very few studies using SDMs explicitly incorporated phenology into SDM estimates, and most of those that did focused on tree species. A few studies have suggested (22) or shown (4, 10, 23, 41) that incorporating the seasonal niche provides better predictions of species’ occurrences than annual niche data. For the annual plant *Mollugo verticillata*, species distribution models built from climate records of herbarium specimen collections by month and year were more accurate in predicting spatial occurrences than those using long-term average climate predictors (4). Seasonal models also better predicted occurrences of invasive mustard *Capsella bursa-pastoris* in the novel range than annual niche models (23). There has been a recent plea to integrate different phenological stages into SDMs even for perennial species because different life stages often exhibit different climatic tolerances (reviewed in (24)). For example, establishment conditions at the seedling stage may be more constrained than those for survival at the adult stage (e.g.,(42)). These studies, and ours, argue that seasonal niche estimation provides a more nuanced understanding of the environmental tolerances of organisms. Here, we show the power of leveraging museum collections with knowledge of phenological cueing mechanisms (such as, in our case, germination cues) and of the field environments occupied by organisms (i.e., field soil data) to understand the evolution of the climate niche.

Climate niche similarities across members of a clade that occupy divergent annual environments may indicate strong evolutionary constraint in the conditions tolerated by these species. Cang and colleagues (43) found slow rates of climate niche evolution in 85 largely perennial pairs of sister species across the grass phylogeny, with very small climate niche shifts across sister species pairs, particularly for temperature, similar to our results. Niche shifts in this study were somewhat greater for precipitation, but still much smaller than rates of projected climate change. In a large survey of 266 phylogroups of plants and animals, rates of climate niche evolution were again much lower than the rate of predicted climate change across collection localities (44). Overall, climate niche evolution seems constrained across a large number of taxa, despite the fact that these studies have been conducted across widely different evolutionary scales.

Life history may play a part in the observed niche evolution. Our study shows that for many annual species in the Streptanthus I clade, the climate niche has remained fairly static with diversification across divergent climatic regions due to changes in the timing of germination and reproduction, and shifts to coarse-soiled habitat. Open habitats *per se* may especially favor annual species that require establishment from seed, and such habitats become rarer in more mesic climates (as suggested in (45)), and may be more common on coarse, well-drained soils (46). However, there are perennial members of this clade that persist aboveground for most or all of the year. Perenniality appears to have evolved multiple times across this clade from annual ancestors, including a subclade of only perennial species (27, 47) that occupy open higher elevation habitats with snow (28, 47), suggesting potential climate niche evolution. Rapid climate niche evolution has been found to be associated with life-history shifts from annuality to perenniality in Montiaceae (48). In the Streptanthus I clade, all of the perennial species are herbaceous, and the vast majority occur at high elevations where the growing season is relatively short (47). Climate niche constraint might still be present in the timing of spring growth and reproduction limited to snow-free days, and thus could also result in less realized climate niche evolution than one might initially expect based on year-round persistence.

Spatial variation in the environment allows species to occupy areas of suitable conditions (refugia) in surrounding unsuitable habitats. We found that *Streptanthus* and *Caulanthus* species that had diversified into cooler, more northerly habitats were growing on coarser soils characterized by less water-holding capacity; these species often occur on isolated coarse-soiled serpentine outcrops (46, 49). We show that soil and microhabitat reflect niche constraints above and beyond temperature and precipitation. Climate water deficit (CWD), a combination of aspect, slope, soil characteristics, precipitation and temperature, best demonstrated how plants track suitable conditions across variable landscapes and latitudes (Figure 2B,C,D, Figure 4), in comparison to classical climate variables of temperature and precipitation. In addition to soil properties, geological aspect is likely also allowing species to track suitable environments. Many of these species are noted to occupy south- or southwest-facing slopes (e.g., six species included here have south-facing slopes in herbarium locality descriptions). Such spatial selectivity in aspect has also been demonstrated to enable niche tracking for tree species growing in northern California. Species partitioned niches within a very small reserve (<2 km) such that species with higher range-wide CWD occupied open, south-facing slopes and microrefugia that were hotter and drier than surrounding habitat (12).

Together, differences in soil texture, topography, precipitation, and temperature determine water availability to plants (best represented here by CWD), which we found to be significantly less variable across specimen-specific time windows than annual or seasonal ones.

Huey and colleagues (50) suggested that individual plants should have broad tolerance ranges because sessile individuals cannot move to avoid stresses; however, our data and those of others (1–4) suggest that many species are subsampling suitable seasonal environments and are constructing relatively constant niches via germination cuing and phenological adjustments, despite their sessile nature. Moreover, traits that facilitate seed dispersal by animal species with specific habitat preferences may also allow plants to track suitable niches through directed dispersal (reviewed in (8, 40)), though this mechanism may not be applicable to most of our species. Our results show that plants are subsampling suitable seasonal environments through a variety of means, including 1) by using environmental cues to time seed germination, 2) by slowing or speeding flowering phenology to reproduce under the best conditions, 3) by using passive or directed dispersal of seed to reach suitable new sites or refugia. All of these mechanisms, and arguably many more, suggest that plants are under selection to ameliorate the conditions under which they, or their offspring, grow, and that they may have more constrained environmental tolerances than originally hypothesized (see also (44)). In other systems, traits that respond to changing environments are both plastic and have an underlying genetic basis that is under selection (e.g., (51)).

Our herbarium data, combined with results from prior experiments, shed light on these species’ relative vulnerability to changing rainfall patterns. Specifically, we predict that desert *Caulanthus* species may be most vulnerable, given that rainfall onset has recently been coming later in the fall in the California Floristic Province, and in smaller amounts (52).

Specimen-specific niches indicate that these southern species have shorter growing seasons that do not extend much into the spring/summer months (Figure 1). Moreover, our herbarium-based growing season estimates are consistent with prior experiments manipulating water availability in this system (Pearse et al. 2020), in which low rainfall-adapted *Caulanthus* species did not substantially extend their growing seasons into spring, even when ample water was provided. Other experiments with this clade further demonstrated that species can speed their phenology in response to later germination onset to make up for lost time, but they cannot fully make up for shortened seasons in fitness (53). Thus we predict later rainfall onset trends will shorten the growing season by delaying germination and will reduce fitness for desert species that have rarely, over the century or so of herbarium records we used, been collected later in the season, even if later rainfall does occur (Figure 2).

Herbarium data have known biases (reviewed in (54)), and we have attempted to address some of these. We added error radius distances for all specimens (as in (23)) and rejected specimens for which location was not recorded accurately enough (our median error radius was less than 0.5 km). We also took care to sample specimens that spanned the geographic range and elevations of the collections and species’ ranges, and used only reproductive specimens. For the purposes of our study, biases that are applied similarly to all species (e.g., tendency to collect near roads) may not have large impacts on our conclusions. One potential bias specific to our study is whether species were collected at comparable phenological stages. For example, if one species tended to be collected relatively more in fruit, our estimate for lifetime climate niche would be more accurate for that species than for species collected at an earlier phenological stage. Because of botanical etiquette and practice, most herbarium specimens are collected in reproductive stages (55), and we used only reproductive specimens; moreover, in a separate study, differences in phenological stage, while reducing accuracy, were not great enough to affect overall conclusions (56). While we attempted to be conservative about our estimates of the growing season conditions that specimens experienced, it is possible that differences in phenological stage could affect our estimates of specimen ‘lived climate.’ We think it is unlikely to change our overall conclusions, however, due to large differences between annual climate patterns and seasonal niches. Our climate estimates rely on our knowledge of germination behavior and senescence, which we have explored experimentally for many of these species in common gardens under ambient temperatures and varying watering regimes (30, 31, 53). That said, herbarium collections of specimens from the field collected solely in fruit would provide greater confidence in the estimates of end of life seasonal climatic conditions.

Despite widely divergent annual climates, we find very similar growing niche conditions, especially temperature and CWD, among the 14 annual species we sampled across this clade of jewelflowers. Our results suggest that annual species in this clade are much more constrained in their climatic tolerances than would be predicted by year-round climate estimates based on their geographic distributions. Annual climate data are typically used in SDMs, but we argue these may overestimate the rate and extent of climate niche evolution that has actually occurred in some groups, including our focal clade that diversified from deserts into cooler regions. Here, knowledge of germination cues from previous experiments greatly enhanced our ability to estimate the start of the growing season climate. Similar data could be gathered for other systems–germination cues for annuals, and possibly leaf out cues for perennials–and could be combined with climate records from herbarium specimen collection sites to estimate specimen-lived climate.

Ultimately, niche constancy achieved through niche construction and establishment in suitable spatial refugia indicates that species in this clade have not adapted to divergent climates, but rather have constructed favorable climate niches that are similar across wide geographic regions through phenological shifts and habitat refugia. Fundamentally narrow climatic tolerances could ultimately result in underappreciated climate vulnerability in these species despite the divergent geographic ranges and annual climates they occupy.

## Materials and Methods

We used herbarium specimens to characterize the climate niche of 14 annual species (*Streptanthus* and *Caulanthus* spp., Figure 1) in the Streptanthus I subclade of the larger Streptanthoid clade (27). Species in this clade share an affinity for open, bare, rocky (including serpentine outcrops), or sandy habitats (46), exhibit variable range sizes, and span nearly 8 degrees of latitude and >3000 m in elevation from desert to montane biomes (Figure 1C; Figure S1). Species in this study require pollination for seed set, but are typically self-compatible (57). We selected these 14 species (which represent about a third of the subclade) based on four criteria: 1) sampling across the phylogeny ensuring representation of main subclades, 2) sampling across the latitudinal and elevational breadth that the clade as a whole occupies, 3) sampling species with enough herbarium collections to obtain an adequate sample size (>40 specimens), and 4) limiting our analysis to annual species. We used only annuals for three reasons: 1) we could more precisely estimate their whole lifetime above-ground growing conditions, 2) a large number of the species in the California Floristic Province are annual (58), and 3) annuals comprise the large majority of species (> 83%) within this subclade.

### Specimen selection and georeferencing

To build our dataset, we used the extensive digital collection of the Consortium of California Herbaria available through their specimen data portal (cch2.org). In coordination with another trait-focused herbarium project (59), we sampled up to 200 imaged specimens per species. When greater than 200 specimens were available for a species, we subsampled across counties to spread our sampling across the species’ geographic range (Figure 1). These efforts resulted in inventories of 52-186 herbarium records per species (average = 141), with specimen years spanning 1898-2016. These species have >90% of their recorded distributions from GBIF in CA, except for *C. cooperi*, which has 82% of observations in CA and 100% in CFP (Table S3). We omitted four specimens that had disjunct coordinates far removed from other observations of these species, as these may represent misidentifications or inaccurate coordinates. We used only specimens with reproductive structures present (buds, flowers, or fruits) so that our sample represents only plants that survived long enough to reach the reproductive stage.

Our approach reconstructs the lived climate of each specimen by estimating the start and end of its growing season (see below) and matching that with climate records from the year and location of collection. Geographic data and locality descriptions for each specimen were downloaded from the Consortium of California Herbaria. We checked coordinates and elevations, and added missing ones using Google Earth, complemented by the USGS Historical Topo Map database (60) and information from google searches of locality descriptions. For each specimen, we added error radius distances (as in (12, 23)), estimated as the distance at which the provided description would no longer be logical. Specimens for which we could not determine coordinates with <=5 km error radius were excluded from analyses (though the large majority of specimens had much more precise coordinates, mean = 1.13 km, median = 0.5 km).

### Estimating the climate conditions experienced by specimens

To quantify the climate conditions experienced by each specimen, we first estimated when the plant was alive above ground, i.e., the duration of time from when the individual germinated to when it was collected. To estimate germination month, we drew on our prior experiments on germination behavior in this clade with many of the same species—all of the annual species in (30) are included here. In this prior work, we conducted staggered experimental waterings of seed cohorts every two weeks under outdoor conditions from mid-September through December to identify preferred germination conditions (30). We found that most (9 of 11) species had very high or greatest germination fractions at early fall rainfall dates, as has also been found in a number of other desert annual species (61). This result is consistent with adaptation to capitalize on the short winter wet season characteristic of Mediterranean climates. Therefore, using historical weather data, we estimated the germination month for each specimen as the first month in the water year (see below) preceding specimen collection when more than 25 mm of rain fell at the specimen collection site. The threshold rainfall quantity selected was based on results of prior studies finding that more than 25 mm of rain was needed to trigger biological activity in plants living in arid environments (62–64), as well as on field observations in this system (6), and greenhouse experiments in which simulated rainfall to saturation resulted in germination of all species (30).

We estimated germination dates at a monthly scale because weather data at a finer temporal grain are not available for the wide range of years in which specimens for this clade were collected. Typically, the water year in the western United States is calculated from October 1-September 30 (e.g. (52)), but we wanted to account for early September rains in some years, so we set our water year from September 1-August 30. We limited plausible germination dates to September through February based on previous studies (6, 30), and to accommodate the possibility of rare late germination events in February. If there were no rain events greater than 25 mm in the months preceding the specimen’s collection, we assumed the month with the greatest rainfall preceding March was the germination month; these criteria eliminated 8 specimens for which there was no detected rainfall event prior to March.

Total specimen growing season was estimated as the observed germination month to the month of collection (Figure 1B). Among the specimens we used, ∼71% had one or more fruits at the time of collection, so we consider the collection date to be an imperfect but acceptable proxy for the end of life in these annual species. For example, a specimen collected on April 25, 1983, would be estimated to start its growing season during the first month when >25 mm of rain fell at that specimen’s geographic location (September 1982), and end its season in April 1983, at the end of the specimen’s collection month. Species-level averages for the end of the season ranged from early April to mid-June (Figure 1B), with 98% of specimens collected in March-July. We extracted monthly values of mean temperature, total precipitation, and total climate water deficit (CWD) from the California Basin Characterization Model with a spatial resolution of 270 m (BCMv65) (65) for each specimen’s location. The Basin Characterization Model estimates CWD based on evapotranspiration, calculated from solar radiation and incorporating aspect (topographic shading), cloudiness, snow and permeability of underlying bedrock and provides historic data beginning in 1896 (66). Because CWD incorporates temperature, precipitation, physiographic properties and soil permeability (65), it is a very comprehensive metric for sessile plants (67, 68) and especially so for this clade that occupies a large variety of elevations and soil types (46).

We estimated the climate niche for each specimen over three different time windows. In each case, we summed precipitation, summed CWD, and averaged temperature over the selected months. First, we summarized annual climate conditions for each specimen using monthly climate data from September in the year prior to collection until August in the year of collection from the collection site. These annual values are typically used in species distribution and ecological niche models to characterize suitable conditions (24, 69); we refer to these estimates as the “annual climate niche”. Second, we estimated the “specimen-specific seasonal climate niche” using climate data from the estimated month of germination of each specimen until that specimen’s collection month; this period circumscribes the lived climate that each specimen experienced. Lastly, we summarized climate data across a fixed seasonal interval based on the average growing season across the whole clade, hereafter the “clade-seasonal niche”. For these estimates we used a single temporal window that represents the average start and end of life for all species in this study across the clade. This average window spans October in the year prior to collection through April in the year of collection and allows us to compare climate niches across a common time window that samples only months in the growing season.

### Characterizing evolution of the climate niche

Contrasting our specimen-specific seasonal niche estimates with climate niches based on annual data allows us to see the extent to which species are sub-sampling from the annual conditions available in their home sites, and allows us to evaluate the extent to which commonly used annual data are effective in characterizing the niche in these species. Comparing our specimen-specific seasonal niches to the clade-seasonal niche allows us to view more precisely how or whether phenological variation among species shapes the climate niche of each species. Specifically, we test whether the variance in the climate conditions that species occupy differs when considering climate during the clade-seasonal time window vs. the specimen-specific time window. If variance is greater among climate averages that are based on specimen-specific time windows, then phenological variation acts to increase divergence in the climate niche. If variance is smaller among climate averages that are based on specimen-specific temporal dynamics, then phenological variation acts to maintain niche constancy across the clade, an indication of phylogenetic niche conservatism. We test for unequal variances between specimen-specific seasonal climate and clade-seasonal climate for temperature, precipitation, and CWD using Levene’s tests implemented in the R package car (70); values were averaged across specimens to obtain one value of each climate variable per species before analysis.

We also evaluated the extent to which climates in the annual, clade-seasonal, and specimen-specific time windows were correlated. Correlation across time windows indicates that species are subsampling climate in time similarly, e.g., that phenological timing traits are not greatly altering the relative climate spaces that these species occupy. Limited correlations across time windows indicates that temporal subsampling substantively changes the relative climate spaces that these species occupy. We quantify the correlations between annual, clade-seasonal, and specimen-specific climate conditions using phylogenetically independent contrasts (pic() function from the ape package (71)) and correlation tests (cor.test() function in the stats package of base R). We then test whether divergence between clade-seasonal and specimen-specific CWD values changes with the average latitude where these species occur—in other words, whether there is a latitudinal pattern in how species temporally subsample climate—using phylogenetic generalized least squares (PGLS) regression (function gls() from the nlme package (72) with a Brownian motion correlation structure calculated using the corBrownian() function in the package ape (71)).

To understand how the seasonal and annual climate niches have evolved in this clade as it expanded and speciated into more northerly regions and more mesic climates, we visualized mean temperature, precipitation and CWD along the phylogeny using the ggtree package (v.3.6.2) (73). To test whether species retained traits of their ancestral niches as they diverged, we evaluated which model(s) of evolution provided the best fit for the evolution of climate niche traits (35) (using the functions bounded_bm() from phytools v.1.9.16 (37) and fitContinuous() from geiger v.2.0.11 (74)). We compare models of 1) Brownian motion (BM), in which climate niche variables evolved as neutral traits, 2) Ornstein-Uhlenbeck (OU), a modified BM model to model constraints on climate niche evolution towards a single optimum, and 3) bounded Brownian motion (BBM): neutral BM evolution with bounded constraints, which themselves could be under selection (75). The limits for bounded Brownian motion model were set as the minimum and maximum values of the climate niche variables. Model comparison was made by comparing AIC values. Models that were within two units of the lowest AIC model were considered to have similar support. We found that Brownian motion and/or bounded Brownian motion were the best models explaining climate niche traits across this clade (Table S2). We tested for phylogenetic signal in climate niche traits by estimating Blomberg’s K also using the phytools package (Blomberg et al. 2003). When we found significant phylogenetic signal in a niche variable (significant K), we also asked whether species’ niches were significantly more constrained than expected based on an unbounded Brownian motion model (K significantly >1) by comparing our estimate of K to a null distribution of K values generated under Brownian motion (38). A pattern in which climate traits were more constrained than expected from a Brownian motion model, would be consistent with phylogenetic niche conservatism (PNC) (Losos 2008; Revell et al. 2008; Cooper et al. 2010). Ultimately, in this study, we think the strongest evidence for niche conservatism is when trait (climate) variation has a smaller range among species for specimen-specific niches than clade-average niches [and is more similar than expected based on phylogenetic relationships, see above].

### Exploring soil microhabitat occupancy across the clade

In addition to temporal selectivity in growing season, the climate that plants experience may be determined by the microhabitats that they occupy, and species may occupy refugia within a matrix of unsuitable habitat. To investigate whether *Streptanthus/Caulanthus* species occupy habitats that subsample the niche space that is broadly available to them, we used soil texture data from two sources. Soil texture affects water holding capacity and is highly variable at fine spatial scales across the landscape (Figure S9) (67, 76). We first extracted soil available water holding capacity estimates for each specimen locality from the California Soil Resource website at 800m^2^ resolution (76), the finest scale available. For 21 of our 1976 localities, water storage data was unavailable and these localities were omitted from this analysis. To estimate the soil niches that are broadly available in these areas, we also extracted soil water holding capacity from 100 random points within a 20 km buffer radius around each locality. In addition, we leveraged field soil data collections from herbarium specimen sites collected in a previous study (46). In that study, we sampled the top 30 cm of soil underneath 1-3 *Streptanthus* or *Caulanthus* plants per site, with an average of 4.3 localities per species. Field soils were sieved into soil fractions and coarse soil fraction was recorded (see (46) for more details). These two data sources have complementary strengths—the gridded data is available for a large number of localities but with imperfect precision and larger spatial resolution, while the field-collected data are highly specific to exact specimen growing sites, but are available for far fewer sites per species.

To ask whether species living in cooler areas are growing in relatively drier microsites that reflect their drier desert origins, we evaluated the relationship between annual temperature and CWD vs. the coarse soil fraction of field-collected samples using two separate PGLS models (function gls() from the nlme package (72) with a Brownian motion correlation structure calculated using the corBrownian() function in the package ape (71)). Both climate data and soil data were averaged across localities to obtain one value per species before analyses. We use the gridded soil data to test whether species are differentially occurring in more water permeable (and likely drier) soils relative to what is available. We calculated the difference between the soil water storage capacity in grid cells where specimens occurred and the water storage capacity available in the surrounding area. We represented the water storage capacity of the area surrounding each occurrence by taking the average of our 100 randomly selected grid cells. We calculated the average difference between occurrences and these random points for each species. We then tested whether these differences were related to the average annual temperature or CWD of each species using two separate phylogenetic generalized linear models as described above. We predicted that if species are retaining elements of the ancestral desert niche, species occupying cooler or more mesic climates might establish in areas that are more high-drainage than the surrounding area, that is, with a more negative soil water storage delta. We also tested whether these deltas were consistent in their direction with a two-tailed sign test using the binom.test() function in the base R stats package. All data wrangling and statistical analyses were conducted in R version 4.4.3 (77).

## Acknowledgments

M. W. Schwartz, A. A. Agrawal, J. T. Anderson, T. M. Givnish, J. Hille Ris Lambers and E. Edwards provided helpful suggestions that improved the manuscript; the content remains our full responsibility.

## Author contributions

SYS and MGB conceptualized the paper and SYS wrote the first draft; all authors contributed to refining ideas and writing. MGB and SJW conducted analyses with input from SYS. MGB spearheaded herbarium data sampling and collection, with help from LL and SYS; NIC collected soil data with SYS. Herbarium work was supported by NSF DEB 1831913 to JRG, JS, JM and SYS with collaborator NIC; soil collections and analyses were funded by NSF DEB 0919559 to SYS and Conacyt Award 187083 to NIC.

## Data availability

All data and code for these analyses will be publicly archived upon manuscript acceptance and will be shared with reviewers upon request.

## Supplementary Figures and Tables

**Figure S1.**
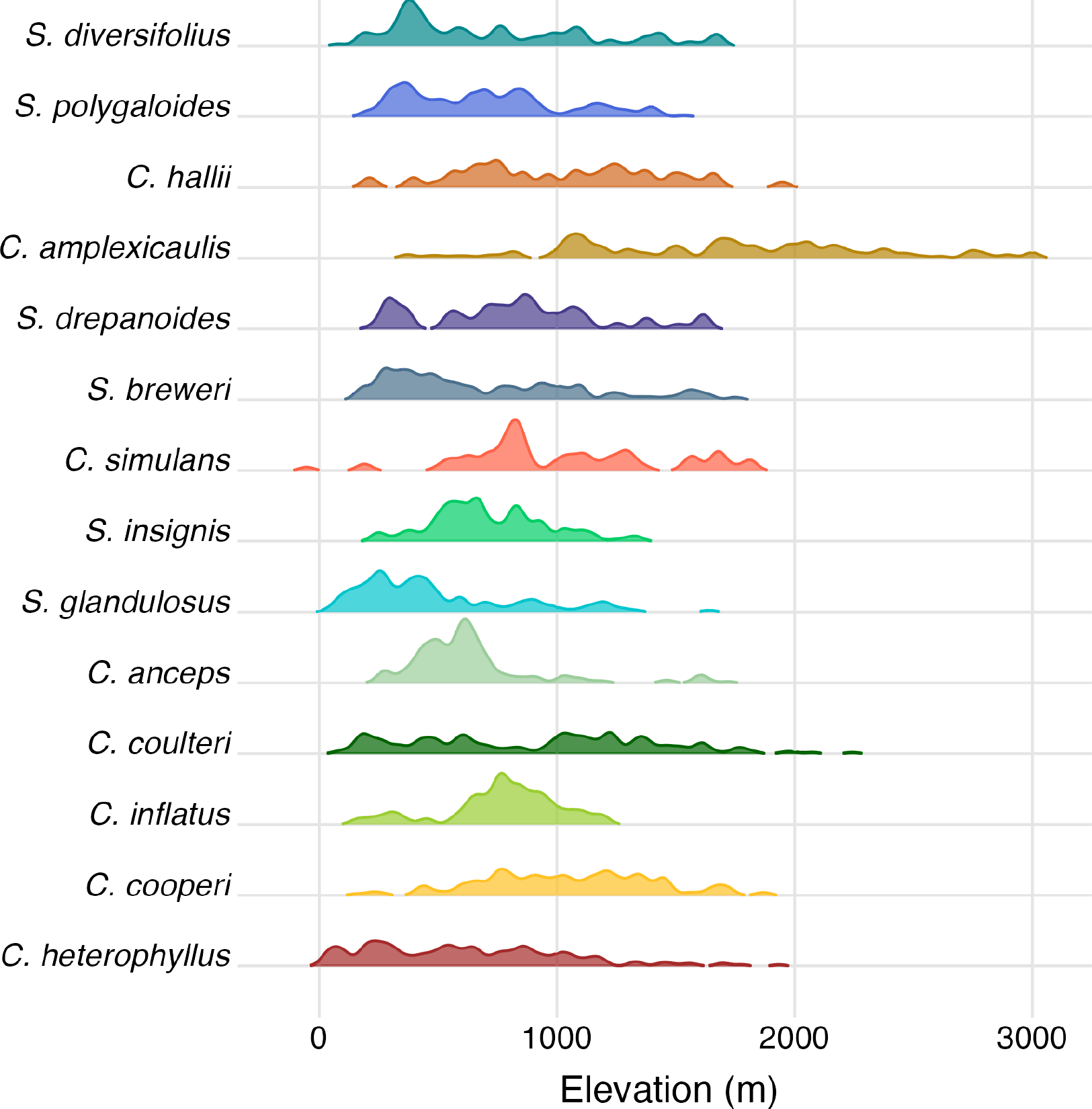
Distributions of specimen elevations (m) for each species. Heights of ridges reflect densities of elevation data scaled within each species. Species are ordered based on the phylogeny plotted in Figure 1.

**Figure S2.**
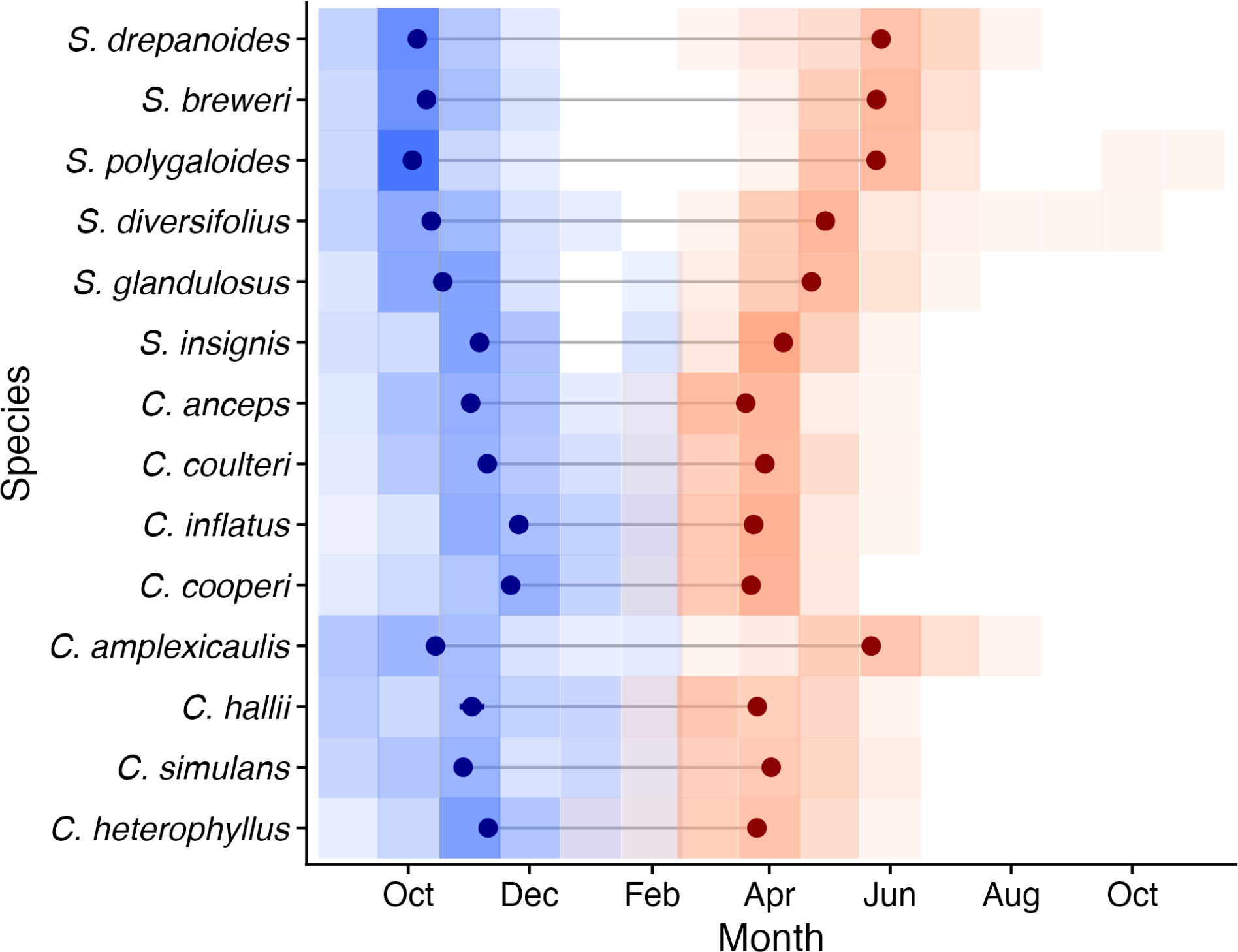
Differences in growing season among the species arranged by latitude (these data are also shown in Figure 1B). Shaded tiles represent the relative proportion of specimens that were estimated to have germinated (blue) or that were collected (red) in a given month for each species. Start month of the growing season was estimated for each specimen as the first month in the fall prior to collection, starting in September, with greater than 25 mm of precipitation. End month of the growing season was the month of collection of each specimen. Specimen-level values were then averaged for species-level estimates of start month and end month, these are depicted with the darker point ranges. *C. amplexicaulis* grows at higher elevations than other species (Figure S1), which may contribute to its deviation from a latitudinal trend in growing season length.

**Figure S3.**
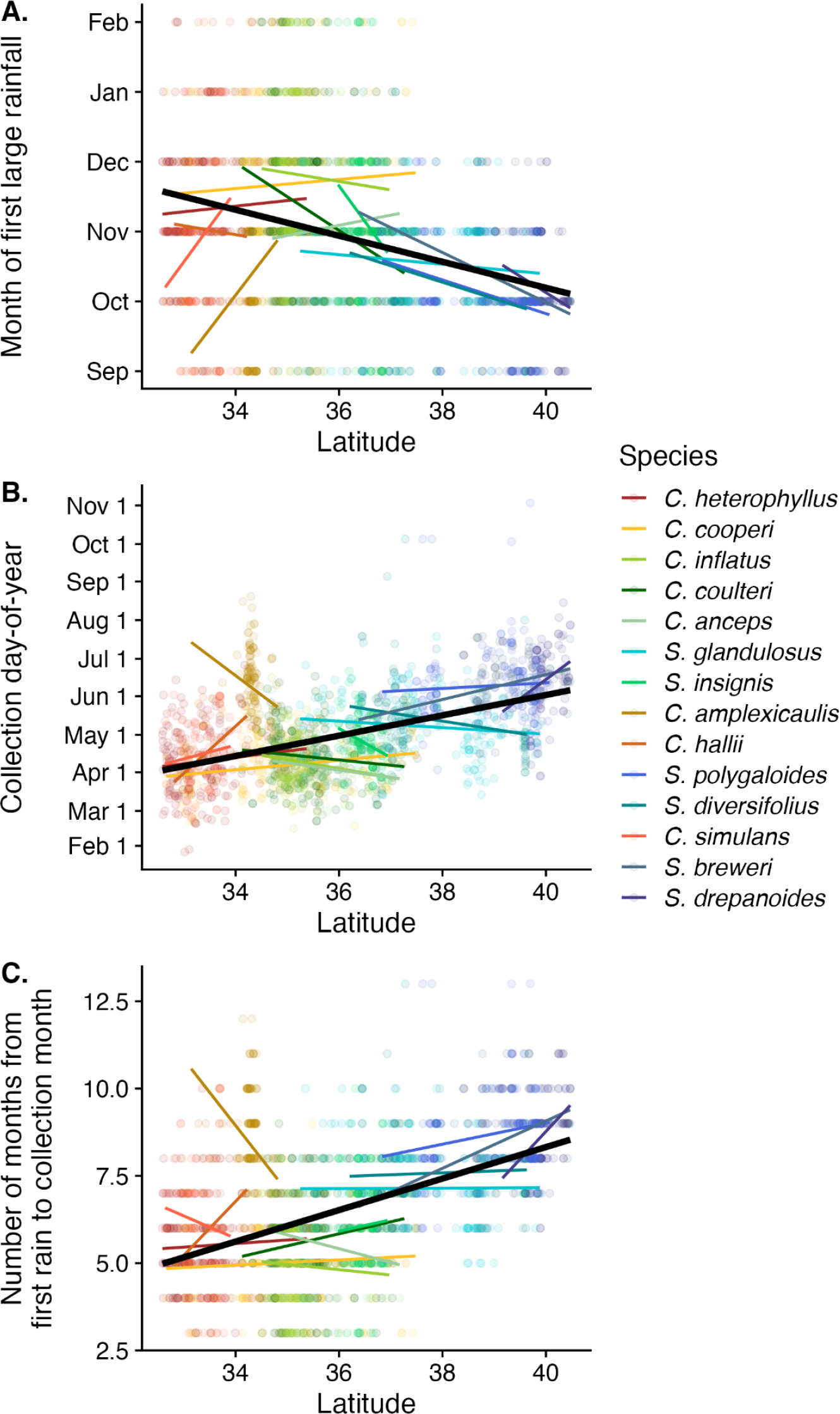
Phenological metrics plotted against latitude. In each panel, trends for individual species are shown with colored lines and the clade-wide pattern is shown with the thick black line. (A.) Species at high latitudes tend to get a first large rain event earlier in the year than those at lower latitudes. (B.) Species at high latitudes tend to be collected later in the year than those at lower latitudes. (C.) Species at high latitudes tend to have longer estimated lifespans than those at low latitudes. Note that trend lines are from linear regressions that do not consider phylogeny and are meant as visual aids rather than statistical estimates.

**Figure S4.**
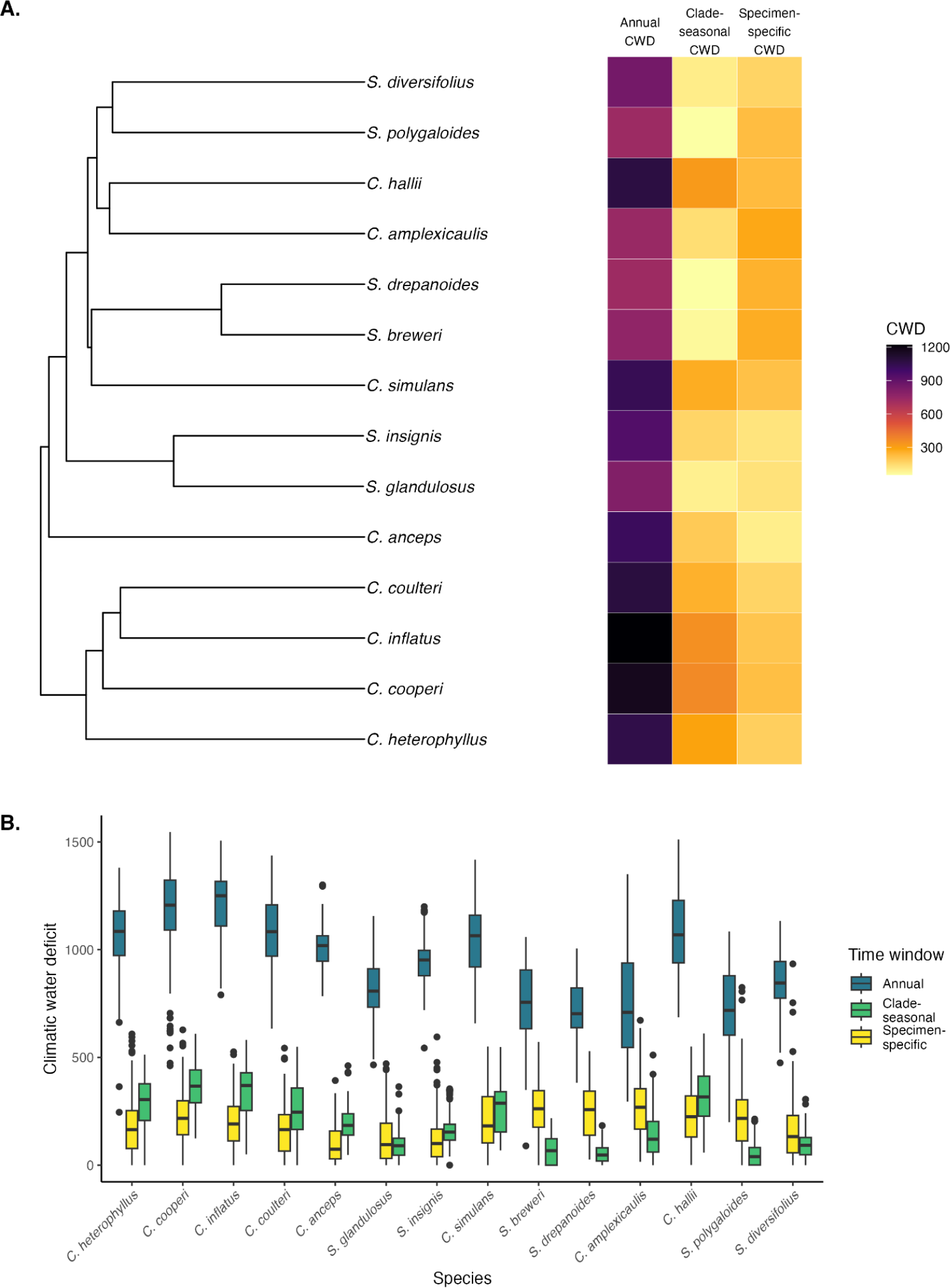
(A.) Climate water deficit (CWD), which estimates water stress from precipitation, temperature, aspect and soil properties) for annual, average clade-wide seasonal, and specimen-specific seasonal time windows plotted alongside the phylogeny. The same color scale is used for all three time windows. (B.) An alternate visualization of the same annual, average clade-wide seasonal, and specimen-specific seasonal CWD; this panel highlights the variation among specimens within species and the magnitude of differences between the three time windows. Species are in the same order as branch tips in (A.)

**Figure S5.**
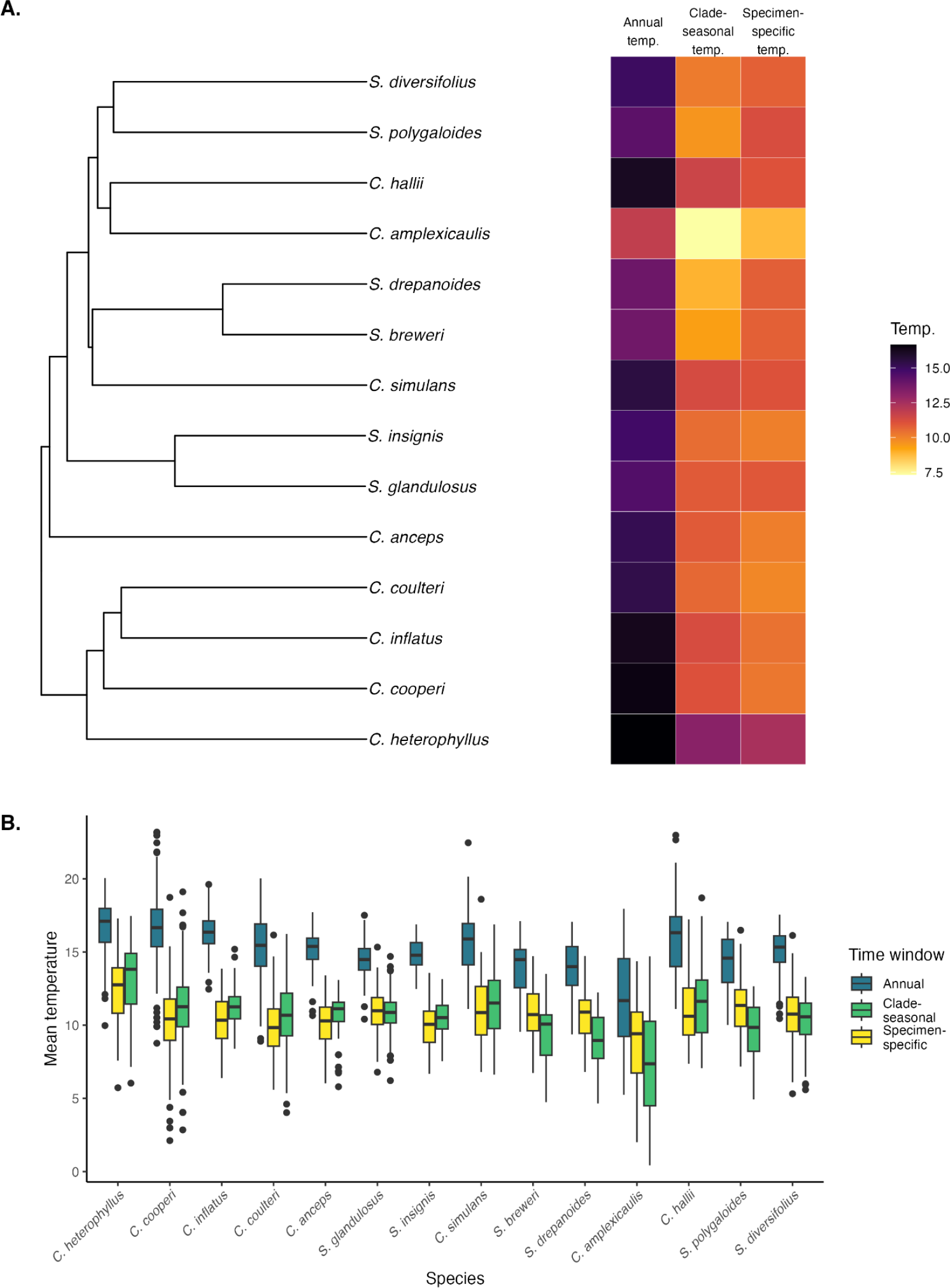
(A.) Average temperature (°C) for the annual, average clade-wide seasonal, and specimen-specific seasonal time windows displayed alongside the phylogeny. The same color scale is used for all time windows to display the differences in average temperature. (B.) An alternate visualization of the same annual, average clade-wide seasonal, and specimen-specific seasonal temperature; this panel highlights the variation among specimens within species and the magnitude of differences between the three time windows. Species are in the same order as branch tips in (A.).

**Figure S6.**
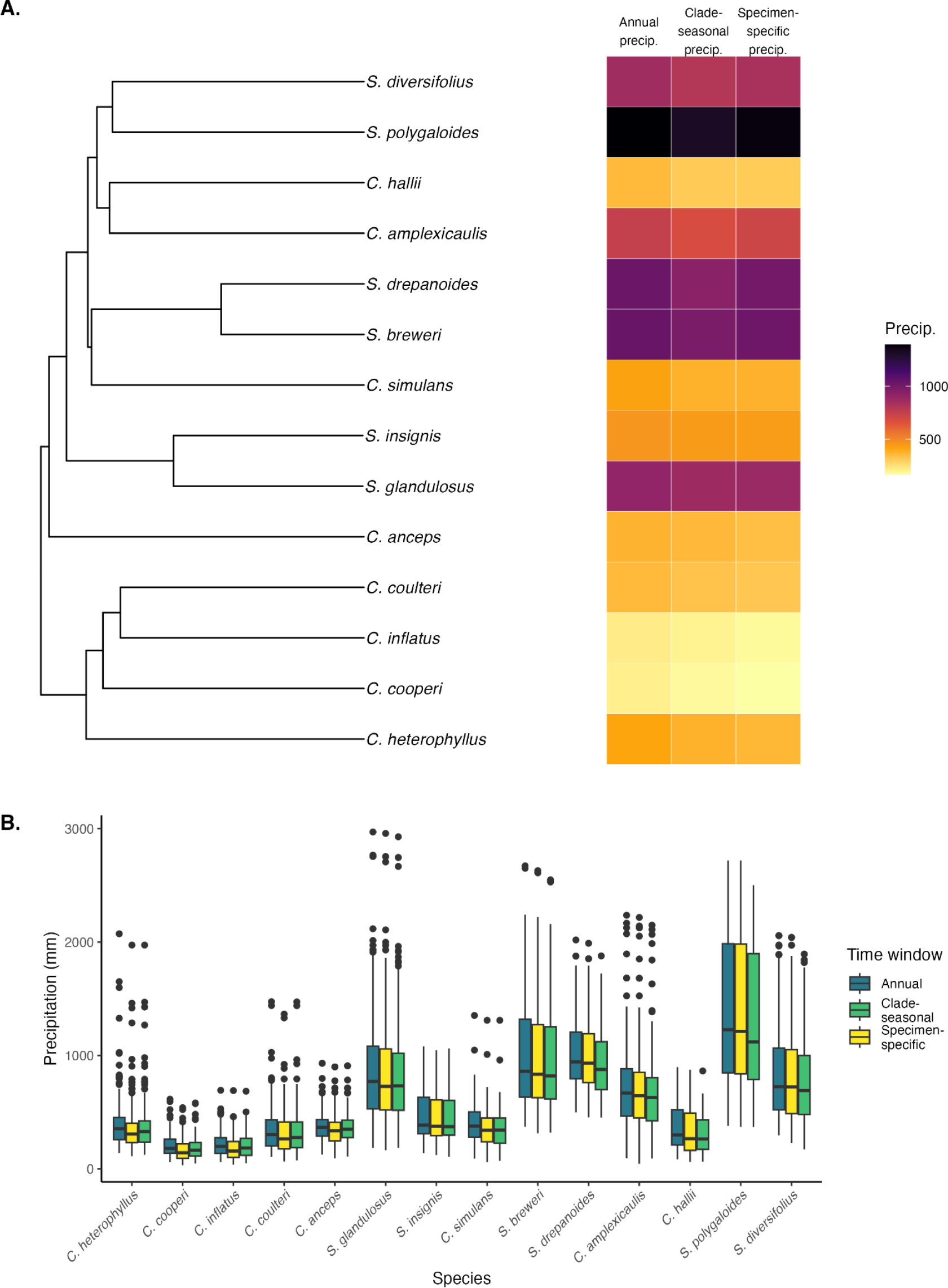
(A.) Summed precipitation (mm) for the annual, average clade-wide seasonal, and specimen-specific seasonal time windows displayed alongside the phylogeny. The same color scale is used for all time windows. (B.) An alternate visualization of the same annual, average clade-wide seasonal, and specimen-specific seasonal precipitation; this panel highlights the variation among specimens within species and the magnitude of differences between the three time windows. Species are in the same order as branch tips in (A.)

**Figure S7.**
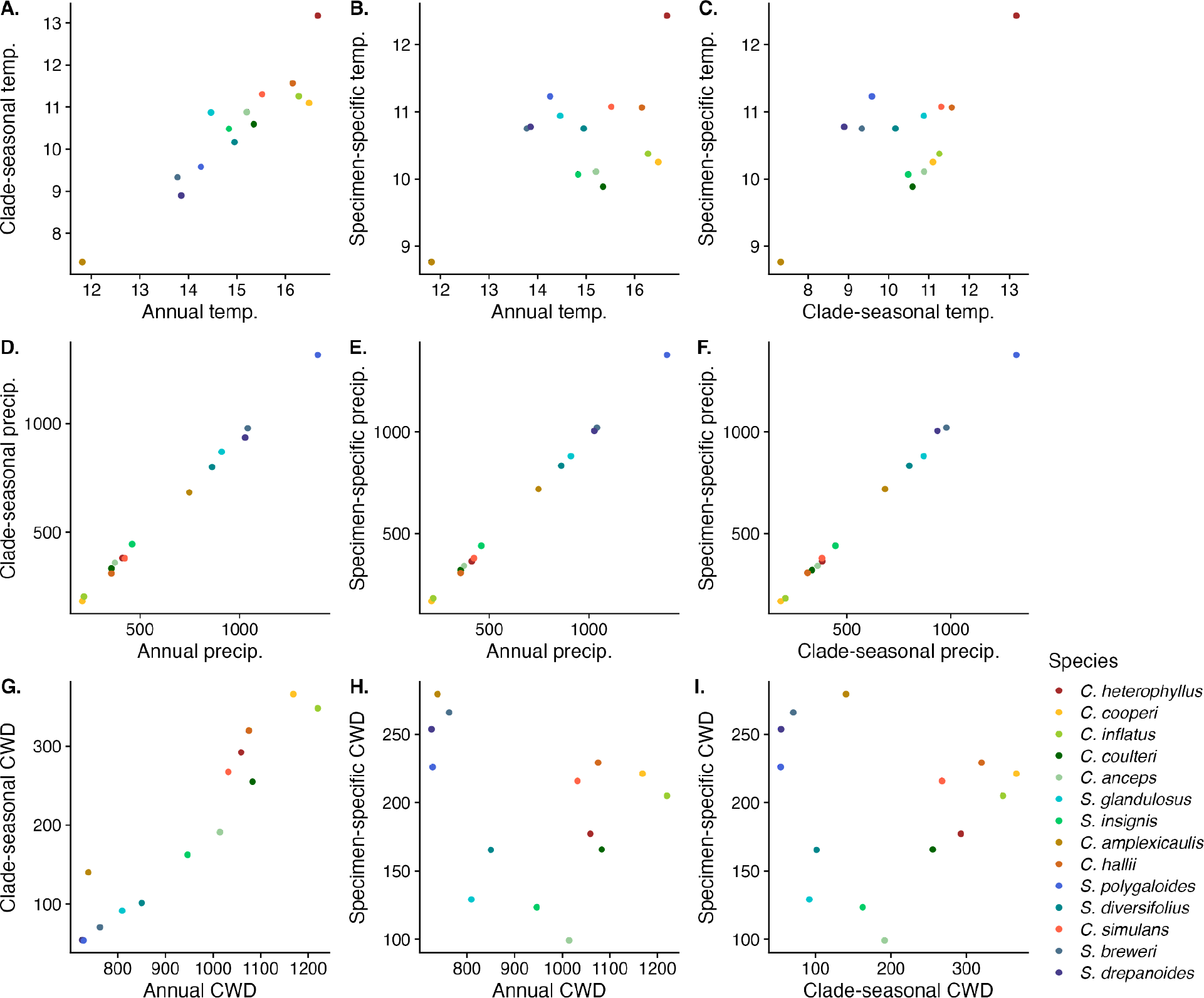
Scatterplots showing relationships over different time scales of average temperature (A.-C.), summed precipitation (D.-E.), and summed climatic water deficit (CWD; G.-I.). Panels are arranged to match the correlation tests in Table 1. (A., D., G.) depict annual vs. clade-seasonal niches. (B., E., H.) depict annual vs. specimen specific niches. (C., F., I.) depict clade-seasonal vs. specimen-specific niches. Correlations are significant in all panels except (H.) and (I.) (see Table 1).

**Figure S8.**
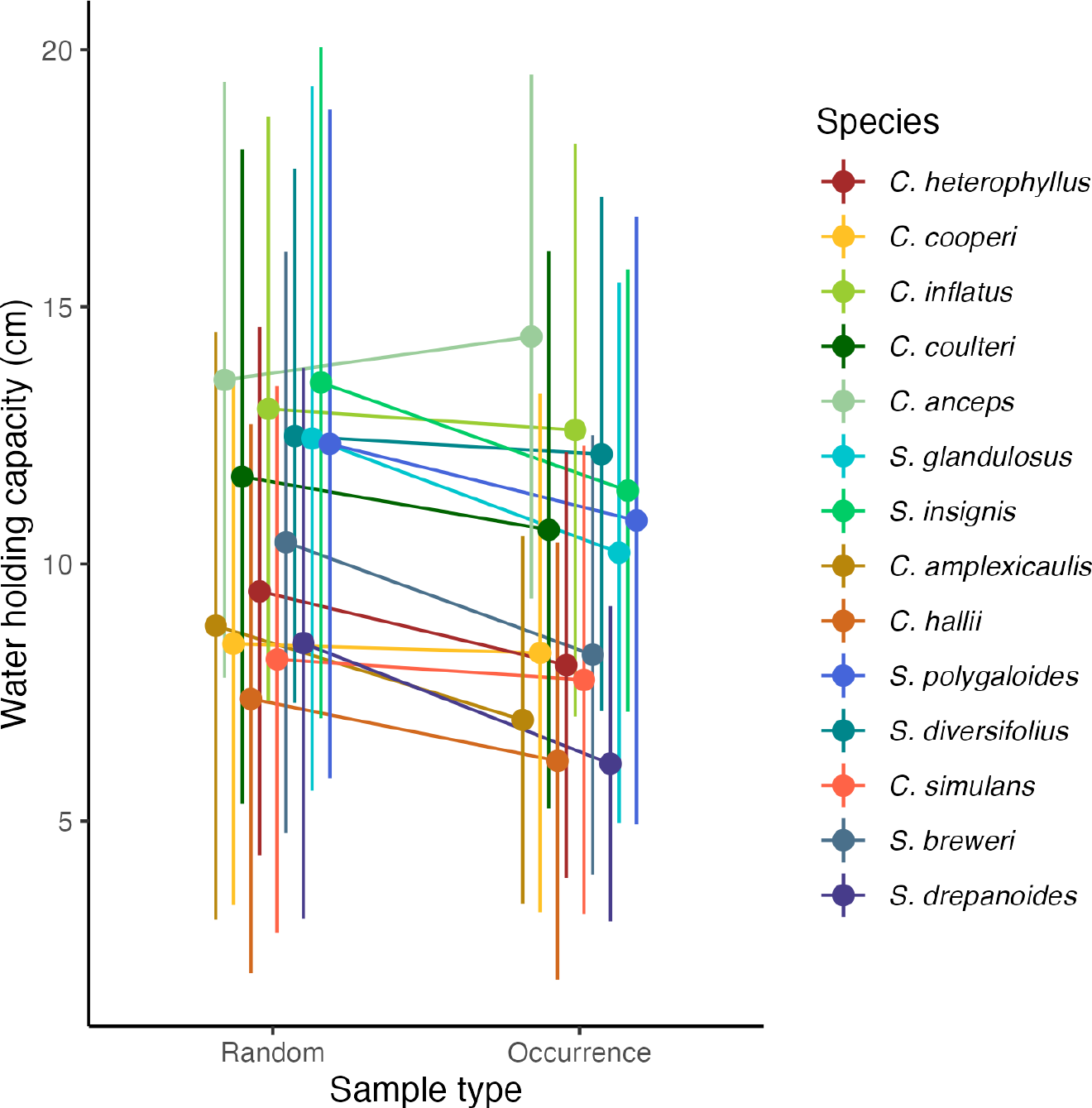
Water holding capacity from occurrence points and from 100 randomly placed points within 20 km of each occurrence. Values were averaged for each species (i.e., averaged across all occurrences for a species, and averaged across the 100 random values x the number of occurrences for each species). Error bars represent standard deviations. Species-averages from occurrence points have lower water holding capacity than the surrounding areas (13/14 species; sign test p = 0.0018).

**Figure S9.**
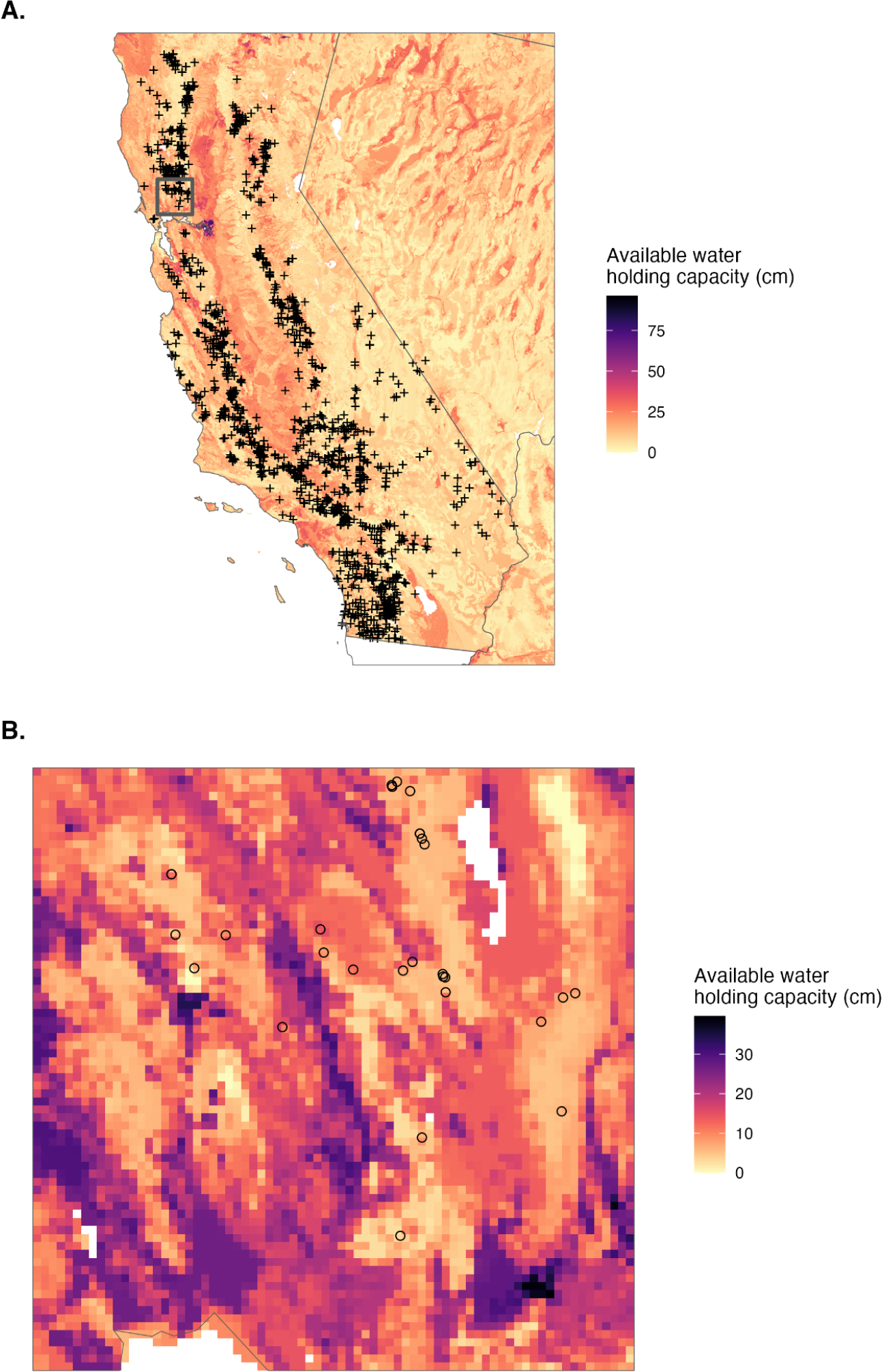
(A.) USDA-NCSS soil survey data within 800 m grid cells illustrating spatial variation in water holding capacity (Walkinshaw *et al*. 2023). Symbols represent locations of specimens used in this study. (B.) Zoomed in view of the area in the grey square in (A.) showing how specimen occurrences in northern parts of the range are in grid cells with lower water holding capacity than the surrounding areas. Note that color scales differ between the two panels.

**Table S1.** Evaluation of phylogenetic signal (Blomberg’s K), for each of the climate variables for each time window. The three climate variables were total precipitation (mm), average temperature (°C), and total climate water deficit (CWD). Climate values used in these analyses were species’ averages of data from each specimen. When K > 1, p_bm_ refers to the significance of the test for whether there is phylogenetic signal consistent with a Brownian motion model of evolutionary change through time (significant if p_bm_ < 0.05). When K > 1 and p_bm_ <0.05, we tested whether K significantly differed from 1; p_gt1_ represents a test for whether K is significantly greater than 1, which indicates that trait evolution is more constrained than expected under Brownian motion (significant if p_gt1_ < 0.05). Note that there is no phylogenetic signal in the specimen-specific temperature niche, in contrast to other climate variables.

|  | Annual climate niche | Average clade-wide seasonal climate niche | Specimen-specific seasonal climate niche |
| --- | --- | --- | --- |
| Total precipitation | K = 1.22<br>**$p_{bm} = 0.0099$**<br>$p_{gt1} = 0.062$ | K = 1.20<br>**$p_{bm} = 0.013$**<br>$p_{gt1} = 0.074$ | K = 1.22<br>**$p_{bm} = 0.010$**<br>$p_{gt1} = 0.055$ |
| Average temperature | K = 1.16<br>**$p_{bm} = 0.017$**<br>$p_{gt1} = 0.11$ | K = 1.12<br>**$p_{bm} = 0.034$**<br>$p_{gt1} = 0.17$ | K = 0.91<br>$p_{bm} = 0.65$ |
| Total CWD | K = 1.35<br>**$p_{bm} = 0.0029$**<br>**$p_{gt1} = 0.017$** | K = 1.30,<br>**$p_{bm} = 0.0052$,**<br>**$p_{gt1} = 0.026$** | K = 1.22<br>**$p_{bm} = 0.013$**<br>$p_{gt1} = 0.060$ |

**Table S2.**
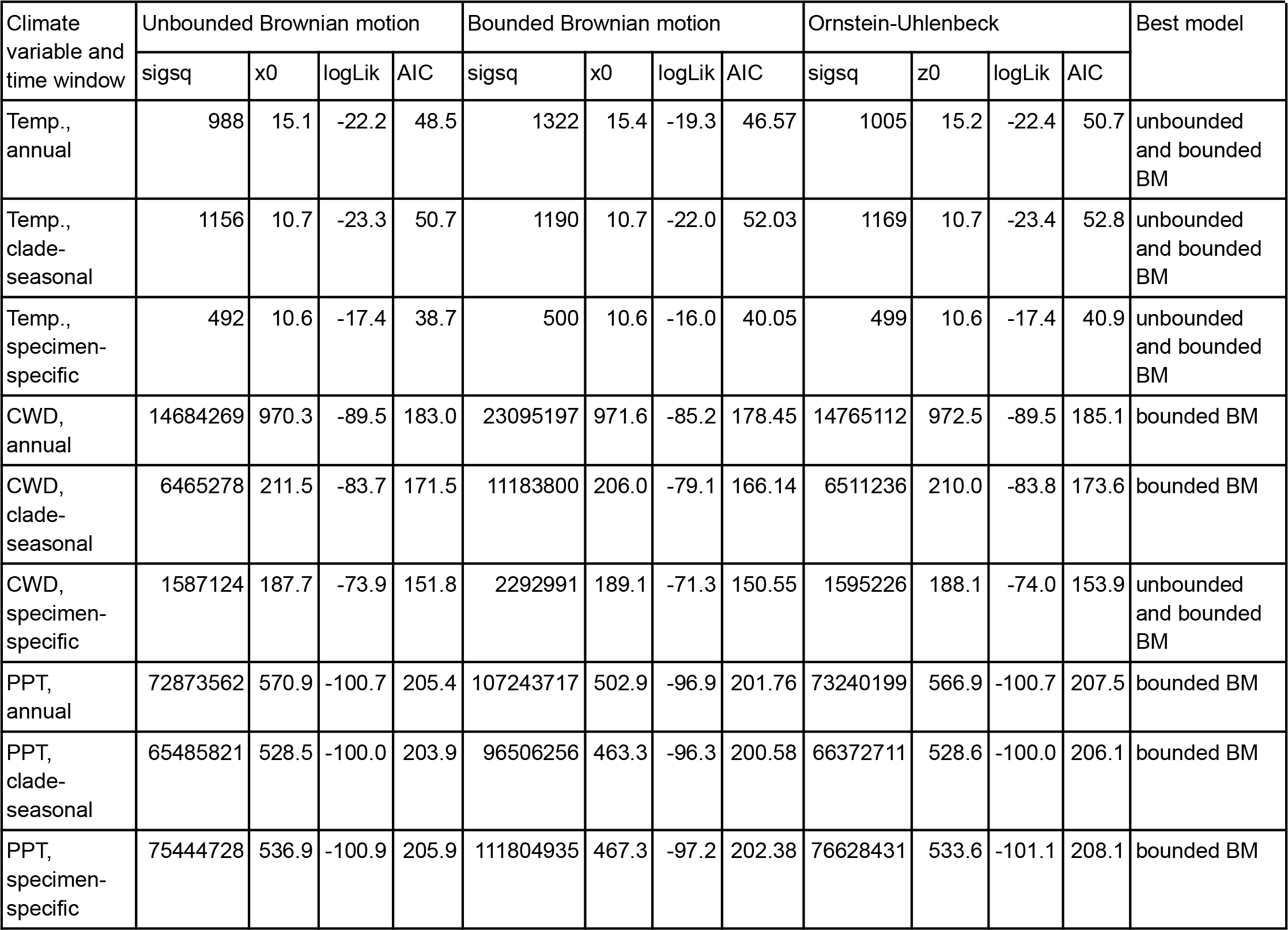
Results of model selection for climate niche evolution. For each combination of climate niche trait and temporal niche, we compared the fit of three models of evolution: 1) Brownian motion (unbounded BM) where climate niche traits evolved as neutral traits as species diverged, 2) Bounded Brownian motion (bounded BM) which is neutral Brownian motion evolution but with bounded constraints, and 3) Ornstein-Uhlenbeck, which is a modified Brownian motion model that constrains climate niche evolution towards a single optimum. Bounded and unbounded Brownian motion models were fit using the phytools package v.1.9.16 (37) and the Ornstein-Uhlenbeck model was fit using the geiger package v.2.0.11 (74). The limits for bounded Brownian motion were set as the minimum and maximum values of the climate variables. Model comparisons were made by comparing AIC values. Evolutionary models that were within two units of the lowest AIC model were considered to have similar support. Reported values from the analyses include the step rate, average amount of change expected in each time step (sigsq), the phylogenetic trait mean or value of the root state (x0 or z0), and the log-likelihood of the models (logLik) (79).

| Climate variable and time window | Unbounded Brownian motion |  |  |  | Bounded Brownian motion |  |  |  | Ornstein-Uhlenbeck |  |  |  | Best model |
| --- | --- | --- | --- | --- | --- | --- | --- | --- | --- | --- | --- | --- | --- |
|  | sigsq | x0 | logLik | AIC | sigsq | x0 | logLik | AIC | sigsq | z0 | logLik | AIC |  |
| Temp., annual | 988 | 15.1 | -22.2 | 48.5 | 1322 | 15.4 | -19.3 | 46.57 | 1005 | 15.2 | -22.4 | 50.7 | unbounded and bounded BM |
| Temp., clade-seasonal | 1156 | 10.7 | -23.3 | 50.7 | 1190 | 10.7 | -22.0 | 52.03 | 1169 | 10.7 | -23.4 | 52.8 | unbounded and bounded BM |
| Temp., specimen-specific | 492 | 10.6 | -17.4 | 38.7 | 500 | 10.6 | -16.0 | 40.05 | 499 | 10.6 | -17.4 | 40.9 | unbounded and bounded BM |
| CWD, annual | 14684269 | 970.3 | -89.5 | 183.0 | 23095197 | 971.6 | -85.2 | 178.45 | 14765112 | 972.5 | -89.5 | 185.1 | bounded BM |
| CWD, clade-seasonal | 6465278 | 211.5 | -83.7 | 171.5 | 11183800 | 206.0 | -79.1 | 166.14 | 6511236 | 210.0 | -83.8 | 173.6 | bounded BM |
| CWD, specimen-specific | 1587124 | 187.7 | -73.9 | 151.8 | 2292991 | 189.1 | -71.3 | 150.55 | 1595226 | 188.1 | -74.0 | 153.9 | unbounded and bounded BM |
| PPT, annual | 72873562 | 570.9 | -100.7 | 205.4 | 107243717 | 502.9 | -96.9 | 201.76 | 73240199 | 566.9 | -100.7 | 207.5 | bounded BM |
| PPT, clade-seasonal | 65485821 | 528.5 | -100.0 | 203.9 | 96506256 | 463.3 | -96.3 | 200.58 | 66372711 | 528.6 | -100.0 | 206.1 | bounded BM |
| PPT, specimen-specific | 75444728 | 536.9 | -100.9 | 205.9 | 111804935 | 467.3 | -97.2 | 202.38 | 76628431 | 533.6 | -101.1 | 208.1 | bounded BM |

**Table S3.**
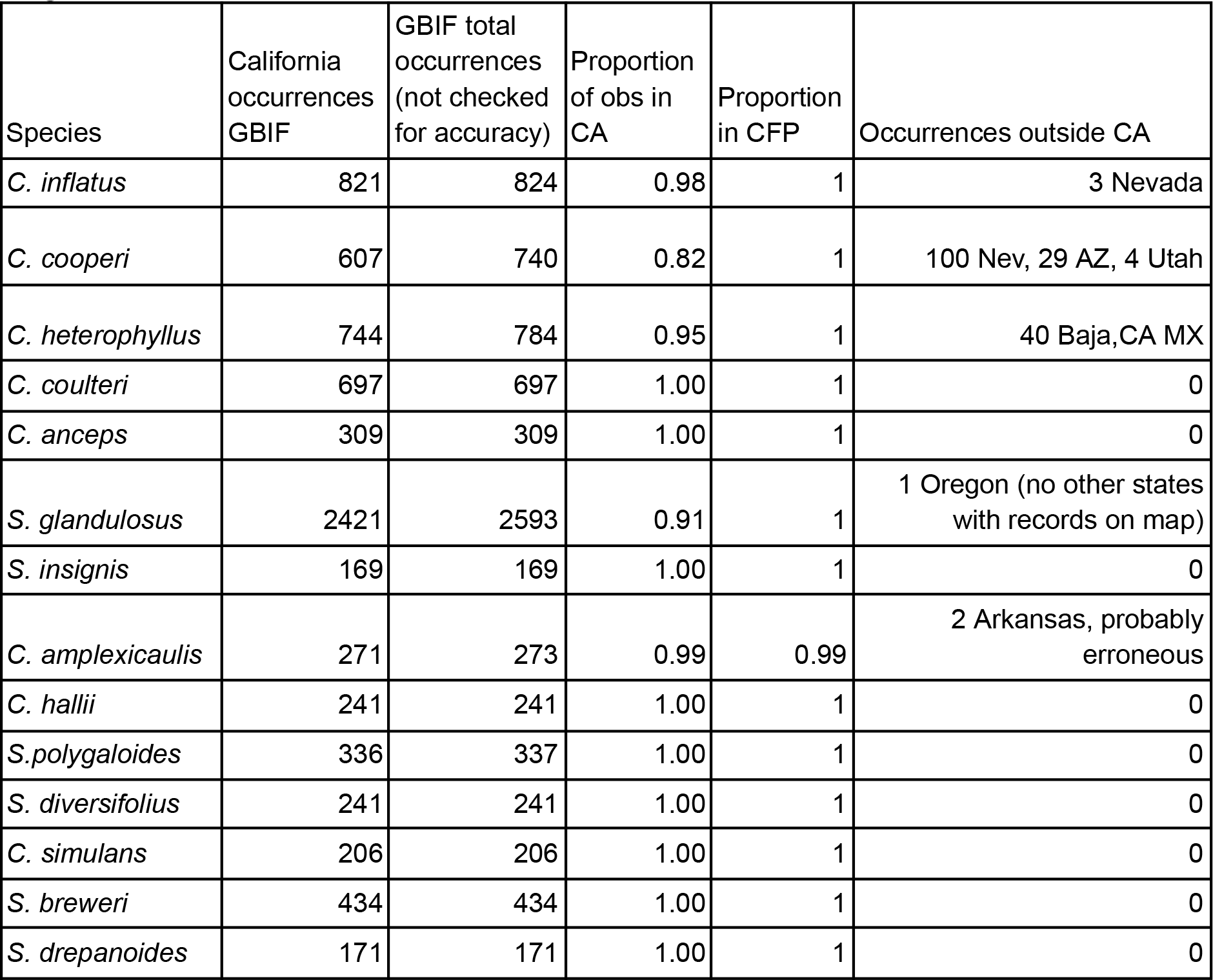
Proportion of species’ occurrences from GBIF in California and the California Floristic Province as of 5/6/ 2025. CA = California. CFP = California Floristic Province, which includes northern Baja California, Mexico, parts of western Nevada and Southern Oregon (Jepson eFlora; https://ucjeps.berkeley.edu/eflora/geography.html). We used the following filters in the GBIF database: Occurrence status: present, Basis of record: herbarium specimen and human observation; Location: coordinates, Continent: North America. GBIF points were not curated for accuracy. In one case (*C. amplexicaulis*), the localities of two observations were disjunct in Arkansas, likely errors in coordinates. All species had >99% of their observations in the California Floristic Province. *C. cooperi* had the lowest proportion of specimens in CA (0.82). Overall, using California-based records provides a good basis for estimating species’ climatic range.

